# Rapid BCMA downmodulation on myeloma cells upon CAR T cell contact is mediated by trogocytosis and BCMA internalization

**DOI:** 10.1101/2022.07.23.501252

**Authors:** Nicolas Camviel, Benita Wolf, Giancarlo Croce, David Gfeller, Vincent Zoete, Caroline Arber

## Abstract

**Background:** Chimeric antigen receptor (CAR) T cell therapy targeting B cell maturation antigen (BCMA) on multiple myeloma (MM) produces fast but not long-lasting responses. Reasons for treatment failure are poorly understood. CARs simultaneously targeting two antigens may represent an alternative. Here, we (1) designed and characterized novel A proliferation inducing ligand (APRIL) based dual-antigen targeting CARs, and (2) investigated mechanisms of resistance to CAR T cells with three different BCMA-binding moieties (APRIL, single-chain-variable-fragment, heavy-chain-only).

**Methods:** Three new APRIL-CARs were designed and characterized. Human APRIL-CAR T cells were evaluated for their cytotoxic function in vitro and in vivo, for their polyfunctionality, immune synapse formation, memory, exhaustion phenotype and tonic signaling activity. To investigate resistance mechanisms, we analyzed BCMA levels and cellular localization and quantified CAR T cell - target cell interactions by live microscopy. Impact on pathway activation and tumor cell proliferation was assessed in vitro and in vivo.

**Results:** APRIL-BB*ζ* CAR T cells in a trimeric ligand binding conformation conferred the best polyfunctionality, immune synapse formation and fast anti-tumor function in vivo in two different mouse xenograft models. Upon CAR T cell – myeloma cell contact, we found rapid BCMA downmodulation on target cells with all three evaluated binding moieties. CAR T cells acquired BCMA on their cell surface by trogocytosis, and BCMA on MM cells was rapidly internalized. Since trogocytosis can lead to CAR T cell exhaustion and presence of CAR T cells does not protect patients from relapse, we investigated whether non-functional CAR T cells play a role in tumor progression. While CAR T cell – MM cell interactions activated BCMA pathway, we did not find enhanced tumor growth in vitro or in vivo.

**Conclusion:** We designed and characterized distinct APRIL-CAR T cells for dual-antigen targeting of MM. Rapid BCMA downmodulation occurred upon CAR T cell – tumor cell encounter independently of whether an APRIL or antibody-based binding moiety was used. BCMA internalization mostly contributed to this effect, but trogocytosis by CAR T cells was also observed. Our study sheds light on the mechanisms underlying CAR T cell failure in MM and can inform the development of improved treatment strategies.

## Introduction

Chimeric antigen receptor (CAR) T cell immunotherapy for multiple myeloma (MM) is currently becoming standard of care in advanced MM.^1 2^ Most approaches currently focus on targeting B cell maturation antigen (BCMA) as a single target because BCMA is critical for maintaining tumor cell phenotypes and expression is limited to the B and plasma cell lineage.^3-8^ While fast and deep anti-tumor responses are consistently reported across clinical trials, the duration of response is quite limited and most patients experience relapse and progression.^9 10^ The reasons for treatment failure are not well understood, but likely involve several mechanisms: (1) antigen escape under immune pressure, (2) insufficient function or exhaustion of circulating CAR T cells, (3) limited CAR T cell migration and persistence, and (4) immune suppression in the MM bone marrow microenvironment. Finally, it is not known whether non-productive binding of persisting exhausted or non-functional CAR T cells to MM target cells could stimulate BCMA pathway activity and contribute to pro-tumorigenic events in residual tumor cells fueling rapid tumor progression. Complete antigen loss has only been described in two patients so far and does not seem to be a frequent event.^9 11 12^

A proliferation inducing ligand (APRIL) is the natural high affinity ligand for two receptors expressed on MM, BCMA and transmembrane activator and CAML interactor (TACI). APRIL-based CARs can overcome BCMA antigen escape in preclinical models,^7 13^ but an early phase clinical trial with a CAR using an APRIL monomer as a ligand binding moiety did not confirm efficacy in patients.^14^ Preclinically, a trimeric APRIL configuration yielded better polyfunctionality and short term anti-tumor function of CAR T cells than monomeric APRIL,^13^ but information on long-term in vivo tumor control and possible mechanisms of resistance remains elusive.

Here, we used molecular modeling and developed and characterized three distinct new APRIL CARs, one monomer- and two trimer-based. We deeply characterized their properties when expressed in human T cells in functional assays in vitro and in vivo. We identified the molecular determinants in monomeric and trimeric APRIL CAR designs that are associated with CAR T cell performance. APRIL CAR T cells with a trimeric binding moiety formed stronger immune synapses and were more polyfunctional than the monomer-based CAR with the BB*ζ* endodomain. In vivo in two myeloma xenograft models, trimer BB*ζ* CAR T cells induced fast anti-tumor responses, but tumors recurred in the follow-up.

Second, we aimed to investigate mechanisms of CAR T cell failure in preclinical MM model systems by comparing a clinically used single chain variable fragment (scFv) based CAR (11D5-3),^3 15^ a human heavy-chain-only (FHVH33) based CAR under clinical development,^16^ and our newly designed APRIL CARs. Upon contact of CAR T cells with myeloma cells, we found rapid BCMA downmodulation on target cells independently of the targeting moiety used. Both CAR T cells and MM cells were actively involved in this process with trogocytosis by CAR T cells and BCMA internalization in MM cells.

Hence, our study provides a better understanding of the function of APRIL CAR T cells and critical insights into determinants of CAR T cell therapy failure in MM when targeting BCMA.

## Methods

### Cell lines

MM.1S (multiple myeloma) cells were purchased from the American Type Culture Collection (ATCC). K562 (chronic myeloid leukemia) and NCI-H929 (multiple myeloma) were purchased from the German Cell Culture Collection (DSMZ). KG-1a (acute myeloid leukemia) cells were a gift from Dr. Stephen Gottschalk, Center for Cell and Gene Therapy, Baylor College of Medicine, Houston, USA. All lines were maintained according to the suppliers’ instructions. Cell lines were authenticated in January 2022 by Microsynth.

### Generation of retroviral vectors

Plasmids encoding for human truncated APRIL (115-250 a.a, Uniprot O75888, excluding the heparan sulfate proteoglycan binding sites of APRIL),^17 18^ full-length BCMA (Uniprot Q02223), full-length TACI (Uniprot O14836), 11D5-3 single chain variable fragment (scFv)^3^ and the human heavy-chain-only FHVH33^16^ were codon-optimized and synthesized by GeneArt (Thermo Fisher Scientific). Retroviral constructs encoding APRIL CARs, 11D5-3 and FHVH33 CARs, BCMA, TACI and BCMA-Emerald were generated using the In-Fusion HD Cloning Kit (Takara, #638933) and expressed in the pSFG retroviral backbone.

### Peripheral blood mononuclear cells from healthy human donors

Buffy coats from de-identified healthy human volunteer blood donors were obtained from the Center of Interregional Blood Transfusion SRK Bern (Bern, Switzerland).

### Generation of CAR T cells and transgenic cell lines

Peripheral blood mononuclear cells (PBMCs) were isolated from buffy coats by density gradient centrifugation (Lymphoprep, StemCell #07851). PBMCs were activated on plates coated with anti-CD3 (1mg/ml, Biolegend, #317347, clone: OKT3) and anti-CD28 (1mg/ml, Biolegend, #302934, clone: CD28.2) antibodies in T cell media (RPMI containing 10% FBS, 2mM L-Glutamine, 1% Penicilin-Streptomycin) with IL-15 and IL-7 (Miltenyi Biotec, 10ng/ml each, #130-095-362 and #130-095-765, respectively) (**Supplementary Table 1**). The day before transduction, a non-tissue culture treated 24-well plate (Grener Bio one, #662102) was coated with retronectin (Takara Bio, #T100B) in PBS (7μg/ml, 1ml per well), and incubated overnight at 4°C. Three days after activation, retronectin was removed and the plate was blocked with T cell medium during 15 min at 37°C. Then, media was removed, and retroviral supernatant was centrifuged onto the retronectin coated plate at 2000g for 1hour at 32°C. Retroviral supernatant was gently removed and activated T cells at 0.15×10^6^ cells/ml was added, and centrifuged at 1000g for 10 min at 21°C. Cells were then incubated at 37°C/ 5% CO_2_ for 3 days. For the generation of transgenic cell lines, cells were counted the day of transduction (0.15×10^6 cells/ml) and performed as described for T cells. After 48 to 72 hours of transduction, T cells were harvested and further expanded in T cell media containing IL-7 and IL-15. In selected experiments, CAR T cells were positively selected with a CD271 selection kit (EasySep, #17849) to enrich for transduced T cells. Transgenic cell lines were then expanded in the appropriate media, checked by flow cytometry for BCMA and/or TACI, or GFP expression, and FACS sorted.

### Sequential co-culture assay

Target cell lines expressing or not GFP-FFLuc were co-cultured with T cells in replicate wells at an effector to target (E:T) ratio of 1:1 (200’000 cells each in a 48-well plate). After three to four days, residual tumor cells and T cells were quantified by FACS combining antibody staining and absolute counting beads (CountBright Beads, Life Technologies #C36950). Fresh tumor cells were added back to replicate wells when >85% of tumor cells were killed at the analysis timepoint. Otherwise, the killing was considered incomplete and T cells were not rechallenged with fresh tumor cells.

### Mouse xenograft experiments

All animal studies were conducted in accordance with a protocol approved by the Veterinary Authority of the Swiss Canton of Vaud and performed in accordance with Swiss ethical guidelines (authorization VD3390). NOD-SCID-*γ*c^-/-^ (NSG) mice were bred and maintained at the animal facility of the University of Lausanne. Animals were 7-11 weeks old at the start of the experiments, both males and females were used, and experimental groups were randomized based on animal weight at start of experiment. Specific information for each model used is provided in the Supplemental Materials.

### Statistical analysis

Descriptive statistics were used to summarize data. For continuous variables, normality distribution was analyzed, and comparisons were made by t-test. When normality testing failed, pair-wise comparison of conditions was performed using Mann-Whitney U-test. Area under the curve (AUC) comparisons were analyzed with unpaired t-test and Welch’s correction (T cell expansion in vitro), or unpaired Mann-Whitney test (non-parametric t-test) (in vivo bioluminescent imaging). Sequential co-cultures and survival of mice were analyzed by Kaplan Meier method and significance assessed with log-rank (Mantel-Cox) test. Multiple comparisons (e.g. BCMA cell surface levels, analysis of sBCMA, T cell phenotypes) were performed by repeated Measures (RM) one-way ANOVA with Tukey’s multiple comparison. Analyses were performed in GraphPad Prism version 8.1.2 or higher. Differences in T cell polyfunctionality was analyzed by dependent t-test for paired samples using Python scipy.stats package (version 1.5.4), where cells co-expressing three or more evaluated markers simultaneously were considered to be polyfunctional. P values <0.05 were considered significant, and the levels of significance are indicated in the Figures and legends.

A detailed description of all methods used is provided in the Supplemental Materials document.

## Results

### APRIL CARs recapitulate physiologic target recognition of BCMA and TACI and can be efficiently expressed in human T cells

To investigate the optimal configuration of APRIL CARs to target MM, we performed molecular modeling of monomer (m) and trimer (t) APRIL CARs in contact with their natural target receptors BCMA and TACI and compared their binding interface to soluble human APRIL (**Figure 1A-B** for BCMA **and Supplementary Figure 1A-B** for TACI). A potentially important role for the histidine residues in position 115 of APRIL is illustrated in the trimer model, as the histidines form close contacts with each other and stabilize the trimer structure around a metal ion (**Figure 1C**). From a structural point of view, the tCAR better recapitulates the physiological target recognition than the mCAR because optimal monomer binding requires binding support of some residues from a second monomer (**Figure 1D**). Absence of a second monomer results in a loss of interaction surface area close to 200 Å^2^ (**Supplementary Text 1**). Moreover, exposing to solvent some non-polar residues normally buried inside binding interfaces could weaken the stability of the mCAR (**Supplementary Text 2 and Supplementary Figure 1C**). Because our modelling suggested a more physiological binding interface for tCARs than for mCARs, we designed the glycine-serine linker between APRIL monomers with a length that accommodates optimal trimer folding. APRIL mCAR and tCAR retroviral vector constructs were generated, including non-signaling control Δ and second generation 28*ζ* and BB*ζ* CARs (**Figure 1E**). Primary human activated T cells were efficiently transduced with all constructs, expanded, and CARs detected on the cell surface (**Figure 1F**).

**Figure 1.**
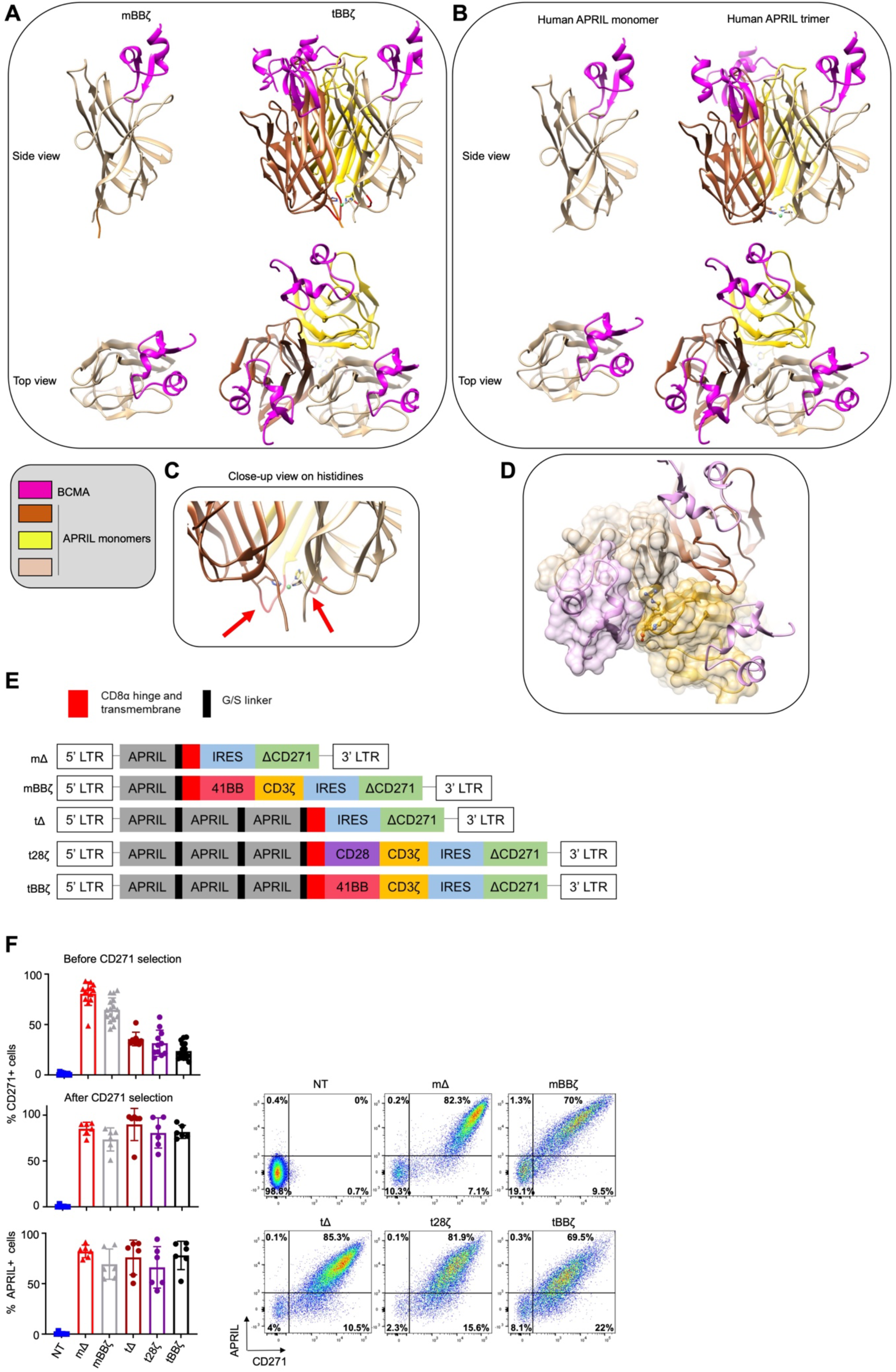
Molecular design and expression of APRIL CARs on human T cells. (**A-D**) Computational models of monomeric (m) and trimeric (t) APRIL ligand binding domains in CAR format (**A**, left: mBB*ζ*, right: tBB*ζ*), compared to human soluble APRIL (**B**, left: monomer, right: trimer) interacting with BCMA (pink). (**C**) Close-up view on histidine residues contributing to the trimer folding, red arrows point to G/S linkers of optimal length. (**D**) Interaction between the APRIL-trimer (each monomer is colored differently in tan, dark brown and goldenrod), and three copies of BCMA (light pink). The surface of one BCMA is shown in transparent pink that is mainly binding one APRIL monomer (surface colored in transparent tan), additional interactions occur with His203, Asp205, Arg206, Tyr208 and His241 residues (in ball and stick representation) of a second APRIL monomer (surface colored in transparent goldenrod). (**E**) CAR construct schemes: mΔ, mBB*ζ*, tΔ, t28*ζ*, tBB*ζ* expressed in the SFG retroviral vector. (**F**) Left: Bar charts represent summary of transduction efficiencies of APRIL CARs in human T cells before or after CD271 selection, based on CD271 marker gene or APRIL cell surface staining (n=8-16 donors; mean±SD). Right: representative FACS plots from one donor, after CD271 selection.

### APRIL CAR T cells are cytotoxic in vitro and in vivo

To probe the cytotoxic activity of the different CAR constructs, we selected several MM cell lines and assessed cell surface BCMA and TACI expression by flow cytometry (**Figure 2A**). NCI-H929 expressed BCMA only, while MM.1S expressed low levels of both BCMA and TACI. The myeloid cell lines KG-1a and K562 were negative for both antigens. We engineered K562 cells to express either antigen alone or in combination and confirmed their cell surface expression levels (**Figure 2A**). To assess anti-tumor function and expansion of APRIL CAR T cells in vitro, we performed a sequential co-culture stress-test. T cells were challenged with fresh tumor cells up to five times, and residual tumor and T cells were quantified 3-4 days after each tumor challenge (**Figures 2B and 2C**). Both mCAR and tCAR T cells produced significant anti-tumor activity against NCI-H929 and MM.1S cells. The mBB*ζ* CAR T cells were the most potent in this assay with the best in vitro T cell expansion capacity (**Figure 2C**). Similar results were obtained when performing the sequential co-culture assay with the engineered K562 targets (**Supplementary Figure 2**). Analysis of the T cell differentiation and activation/ exhaustion phenotype revealed that CD4+ t28*ζ* CAR T cells differentiated faster to an effector memory (T_EM_) state compared to mBB*ζ* and tBB*ζ* CAR T cells even at baseline in the absence of target antigen, while CD8+ mBB*ζ* CAR T cells were more enriched in central memory (T_CM_) phenotype cells compared to control (**Supplementary Figure 3A**). In addition, LAG3 levels were significantly higher on all except tBB*ζ* CD4+ CAR T cells compared to ΔCARs and on mBB*ζ* CD8+ CAR T cells at baseline. PD1 and TIM3 levels were comparable among groups (**Supplementary Figure 3B**). APRIL CAR T cells did not spontaneously secrete significant amounts of IL-2 and did not proliferate in the absence of target antigen (**Supplementary Figure 4**). Overall, these observations indicate absent to low tonic signaling activity of APRIL CARs which was least evident with the tBB*ζ* CAR. Lastly, we evaluated the in vivo anti-tumor function of APRIL CAR T cells in a mouse xenograft model with NCI-H929 cells (**Figure 2D**). mBB*ζ*, t28*ζ* and tBB*ζ* CAR T cells significantly delayed tumor growth (mΔ vs mBB*ζ*: p=0.02, mΔ vs t28*ζ*: p=0.008, mΔ vs tBB*ζ*: p=0.008, n=5) (**Figures 2E-G**), but only tBB*ζ* CAR T cells significantly enhanced overall survival of mice (mΔ vs tBB*ζ*: p=0.003, mBB*ζ* vs tBB*ζ*: p= 0.01, n=5) (**Figure 2H**).

**Figure 2.**
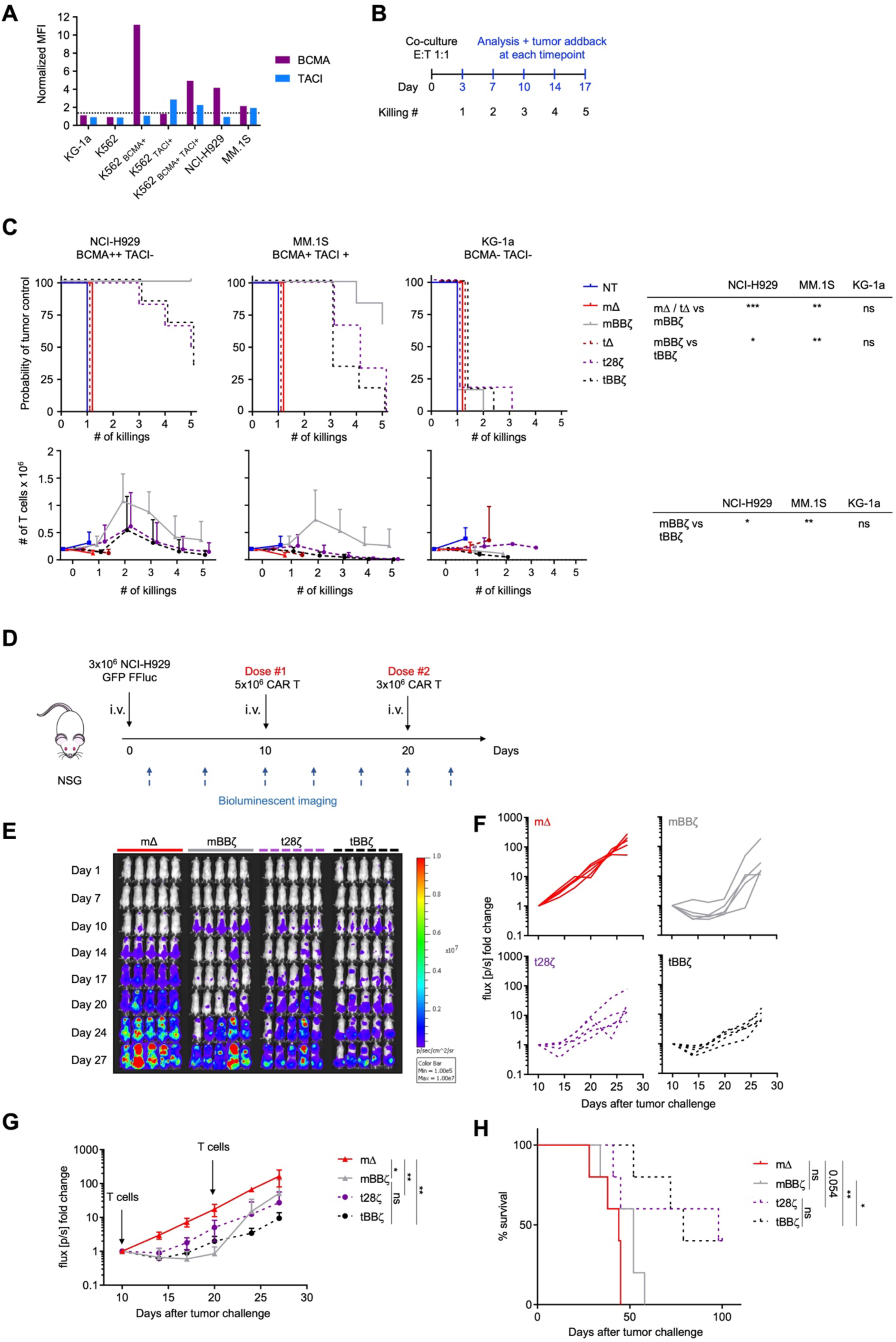
In vitro and in vivo anti-tumor function of APRIL CAR T cells. (**A**) BCMA and TACI expression on cell lines relative to isotype control staining. (**B**) Scheme of rechallenge co-culture stress-test, effector to target ratio E:T 1:1. (**C**) Kaplan Meier curves in the top row depict probability of tumor control in the rechallenge co-culture stress test, n=6 donors, log-rank test (table top right). Line graphs show T cell expansion in co-cultures, n=6, mean±SD, unpaired t-test with Welch’s correction on AUC (table bottom right). (**D**) Scheme of the in vivo experimental timeline. 1 of 2 representative experiments shown, n=5 mice per group per experiment. (**E**) Tumor growth measured by BLI, individual mouse pictures (color scale min 1×10^5^, max 1×10^7^ p/sec/cm^2^/sr). Dorsal view identifies tumor growth in the spine and skull. (**F**) Individual line graph of tumor growth for each mouse per group. BLI flux [p/s] normalized to signal intensity measured on Day 10. (**G**) BLI flux summary n=5, mean±SD. Unpaired Mann-Whitney-U test (non-parametric t-test) on AUC. (**H**) Survival of mice as defined by humane end-point criteria. Log-rank test. (**C, G, H**) Definition of significance levels: ns=not significant, *p<0.05, **p<0.01, ***p<0.001.

### Better T cell polyfunctionality and immune synapse formation is achieved with tBB*ζ* than with mBB*ζ* CARs

By intracellular cytokine staining (**Figure 3A**) and analyses of the immune synapse (**Figures 3B-E**) we assessed CAR T cell polyfunctionality and the strength of T cell – target cell interaction. In both CD4+ and CD8+ CAR T cells, polyfunctionality was enhanced when comparing mBB*ζ* to tBB*ζ* CARs (**Figure 3A and Supplementary Figure 5**) despite considerable donor variability. To further investigate the differences between mBB*ζ* and tBB*ζ* constructs, we performed confocal microscopy and analyzed features of the engineered immune synapse (**Figure 3B**). While actin accumulation at the T cell – target cell interface was comparable between mBB*ζ* and tBB*ζ* CAR T cells, the microtubule organization center (MTOC) localized significantly closer to the synapse in tBB*ζ* compared to mBB*ζ* CAR T cells (**Figures 3C-E**), suggesting the formation of stronger immune synapses with the tBB*ζ* CAR.

**Figure 3.**
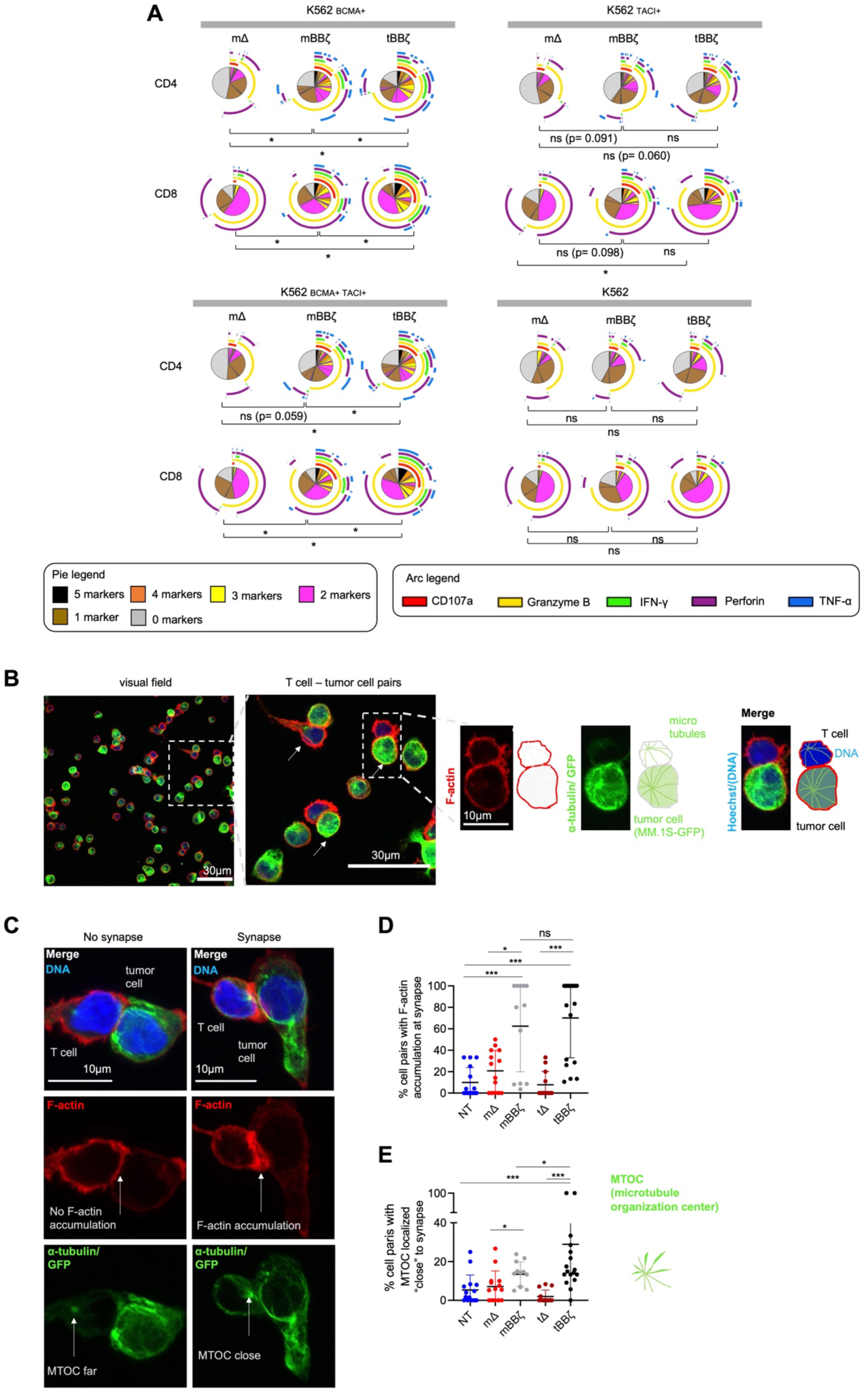
Analysis of polyfunctionality and immune synapses of APRIL CAR T cells. (**A**) Polyfunctionality by intracellular cytokine staining. Pies depict all different combinations of populations expressing 5, 4, 3, 2, 1 or 0 markers, and arcs around pies indicate the analyte detected. Pies represent mean of n=6 donors. Dependent t-test for paired samples was used to compare populations of CAR T cells expressing at least 3 markers. (**B-E**) Fixed cell confocal microscopy for GFP (green, tumor cell), *α*-tubulin (green, showing the microtubule organizing center (MTOC), centrosome), F-actin (red) and DNA (blue, Hoechst). (**B**) Left panel: Representative confocal microscopy field of view showing NT T cells and MM.1S-GFP-FFLuc tumor cells, scale bar 30µm. Middle panel: enlarged inset of left panel, arrows marking T cell - tumor cell pairs, scale bar 30µm, Inset: enlarged view of one T cell (upper cell) - tumor cell (lower cell) pair, scale bar 10µm. Scheme on the right explains the immunofluorescence. (**C**) Left column: representative confocal microscopy images for T cell - tumor cell pair without features of an immunological synapse. Right column: representative images with features of an immunological synapse. First row: Merged images. Second row: F-actin, left: no F- actin accumulation at the cell-cell interface, right: F-actin accumulation at the synapse. Third row: alpha tubulin and GFP. Arrows are pointing towards the centrosome, the major microtubule organizing center (MTOC). Scale bars 10µm. (**D**) Quantification of % cell pairs with F-actin accumulation at the cell-cell interface. Each data point corresponds to one field of view, mean±SD, Mann-Whitney-U test. (**E**) Quantification of % cell pairs with the centrosome close to the cell-cell interface. Each data point corresponds to one field of view, mean±SD, Mann-Whitney-U test. (**A, D, E**) Definition of significance levels: ns=not significant, *p<0.05, ***p<0.001.

### CAR T cells mediate BCMA downmodulation on target cells by trogocytosis and internalization

We next investigated possible mechanisms of resistance when MM cells are exposed to CAR T cells, investigating APRIL CARs and two different well-studied BCMA-targeting CARs (FHVH33 and 11D5-3) (**Figure 4A**).^3 9 16 19^ First, we analyzed BCMA levels on target cells upon contact with the different CAR T cells. MM.1S cells expressed low levels of BCMA that remained stable in the presence of mΔ and tΔ CARs, and decreased significantly upon contact with mBB*ζ*, tBB*ζ* as well as FHVH33Δ, 11D5-3Δ, FHVH33BB*ζ* and 11D5-3BB*ζ* CAR T cells (**Figure 4B**). BCMA reduction on target cells was not due to an increased BCMA shedding as soluble BCMA (sBCMA) levels in the culture supernatant were also reduced in the presence of mBB*ζ*, tBB*ζ*, FHVH33Δ and 11D5-3Δ CAR T cells (**Figure 4C**). However, we detected an increase of BCMA on the cell surface of mBB*ζ* and tBB*ζ* CAR T cells compared to mΔ and tΔ (**Figure 4D**), a process known as trogocytosis.^20^ NCI-H929 cells expressed high levels of BCMA on the cell surface that rapidly decreased in the presence of FHVH33Δ and 11D5-3Δ CAR T cells, but BCMA levels were restored after 24 hours with all ΔCARs (**Figure 4E**). Similar to MM.1S cells, NCI-H929 cells did not downmodulate cell surface BCMA levels upon contact with mΔ and tΔ CAR T cells. However, we detected significant BCMA reduction on residual tumor cells within 6 and 24 hours of co-culture with all BB*ζ* CAR T cells and MM.1S cells, but only at 24 hours with NCI-H929 cells (**Figures 4B and 4E, and Supplementary Figure 6A**). sBCMA levels were decreased with both cell lines (**Figures 4C and 4F**). We also detected BCMA uptake from NCI-H929 cells by mBB*ζ* and tBB*ζ* CAR T cells consistent with trogocytosis (**Figure 4G and Supplementary Figure 6B**). In addition, FHVH33Δ and 11D5-3Δ CAR T cells had high levels of BCMA on their cell surface upon co-culture with NCI-H929 (**Figure 4G and Supplementary Figure 6B**) that were higher than in MM.1S co-cultures (**Figure 4D**). This may be explained by the overall much higher levels of BCMA on NCI-H929 compared to MM.1S. Since treatment with *γ*-secretase inhibitors (GSI) has been shown by others to be a potential strategy to increase BCMA cell surface levels on target cells by reduction of BCMA shedding,^21^ we investigated addition of the GSI LY3039478 during co-culture. When LY3039478 was added at initiation of co-culture, LY3039478 treatment could not prevent BCMA downmodulation in both MM.1S and NCI-H929 cells (**Figures 4H and 4I, and Supplementary Figure 6C**). These results led us to explore BCMA internalization as a potential explanation for reduced BCMA cell surface levels during co-culture. We encoded emerald-tagged BCMA into a retroviral vector (**Figure 4J**) and generated MM.1S-and K562-BCMA-Emerald transgenic cell lines. Co-localization of BCMA-Emerald with cell surface BCMA staining was analyzed by ImageStream upon co-culture with T cells. Contact of 11D5-3Δ CAR T cells with MM.1S cells led to a significant reduction of BCMA co-localization on the tumor cell membrane (NT vs 11D5-3Δ: 66.5±6.3% vs 35.0±16.3%, n=5, mean±SD, p=0.005), with a consequent increase in the fraction of cells with no co-localization (NT vs 11D5-3Δ: 14.2±4.0% vs 44.5±14.1%, n=5, mean±SD, p=0.01) (**Figures 4K and 4L, and Supplementary Figure 7A**). Both K562-BCMA-Emerald and MM.1S-BCMA-Emerald cells co-cultured with 11D5-3Δ CAR T cells significantly downmodulated cell surface BCMA levels, while emerald was only slightly reduced (**Supplementary Figures 7B-C and 7F-G**). 11D5-3Δ CAR T cells gained emerald on their surface (**Supplementary Figures 7D-E and 7H-I**) consistent with trogocytosis. Upon co-culture with APRIL CAR T cells, we found that only tBB*ζ* but not tΔ CAR T cells induced BCMA internalization in MM.1S-BCMA-Emerald cells (**Figures 4M, 4N and Supplementary Figure 8A**) or in K562-BCMA-Emerald cells (**Supplementary Figure 8B**). Very low but significant levels of emerald were also detected on tΔ or tBB*ζ* APRIL CAR T cells consistent with trogocytosis (**Supplementary Figures 8C, 8D and Supplementary Figure 9**). Altogether, these data suggest that BCMA downmodulation on MM cells is mediated by a combination of trogocytosis and BCMA internalization upon CAR T cell exposure with internalization accounting for most of the effect.

**Figure 4.**
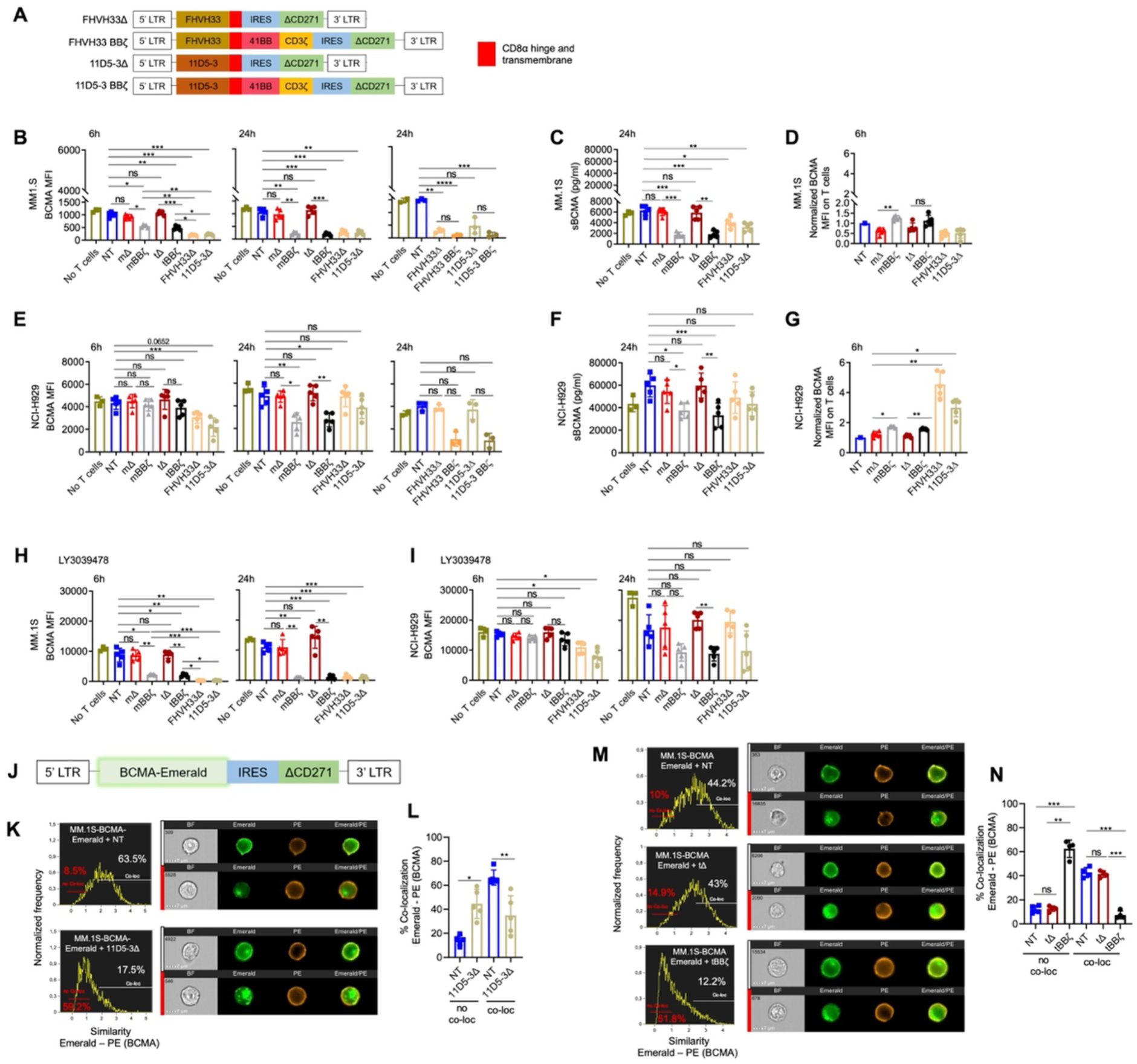
BCMA downmodulation on MM cells by CAR T cell trogocytosis and BCMA internalization. (**A**) Schemes of FHVH33 and 11D5-3 CAR retroviral vector constructs. (**B**) BCMA cell surface levels on MM.1S cells co-cultured with CAR T cells (E:T 1:1) at indicated time-points. (**C**) Soluble BCMA (sBCMA) levels in MM.1S co-culture supernatant (24h). (**D**) BCMA levels on CAR T cells co-cultured with MM.1S cells, normalized to NT (6h). (**E**) BCMA cell surface levels on NCI-H929 cells co-cultured with CAR T cells (E:T 1:1) at indicated time-points. (**F**) sBCMA levels in NCI-H929 co-culture supernatant (24h). (**G**) BCMA levels on CAR T cells co-cultured with NCI-H929 cells, normalized to NT (6h). (**H-I**) Cell surface BCMA levels on MM cells co-cultured with CAR T cells (E:T 1:1) at indicated time-points, in the presence of *γ*-secretase inhibitor, 0.1μM Crenigacestat (LY3039478). n=5, mean±SD, One-way ANOVA with Tukey’s comparison. (**J**) Scheme of BCMA-Emerald retroviral vector with ΔCD271 as selectable marker. (**K-N**) MM.1S-BCMA-Emerald cells were co-cultured for 1h with NT or 11D5-3Δ CAR T cells (**K-L**) or with NT, tΔ or tBB*ζ* CAR T cells (**M-N**). Co-localization of BCMA-Emerald and BCMA cell surface staining (PE) was analyzed by Image Stream. (**K, M**) Representative images and gating. (**L, N**) Summary graphs show percentage of no co-localization (no co-loc) or co-localization (co-loc) of BCMA-Emerald and cell surface BCMA (PE) in MM.1S-BCMA-Emerald cells. n=4-5, mean±SD, paired t-test. (**B-I, L, N**) Definition of significance levels: ns=not significant, *p<0.05, **p<0.01, ***p<0.001, ****p<0.0001.

### CAR T cell - tumor cell interactions trigger BCMA signaling in MM cells

Since disease relapse with BCMA positive disease was reported in clinical trials despite CAR T cell persistence,^9 22^ we next assessed if BCMA directed CAR T cells have the potential to activate BCMA signaling in MM cells. Upon short co-culture of CAR T cells with MM.1S and NCI-H929 cells, mΔ and tΔ CAR T cells induced only minimal NFκB activation while mBB*ζ*, tBB*ζ*, FHVH33Δ, FHVH33-BB*ζ*, 11D5-3Δ and 11D5-3 BB*ζ* CAR T cells all induced strong NFκB activation (**Figure 5A**). To investigate if differences in T cell – tumor cell interactions could explain differences in NFkB pathway activation in MM.1S cells, we next investigated target cell killing and the quality of the interaction (immune synapse formation, loose interactions, and their durations) at the single cell level by live time-lapse light sheet microscopy (**Figure 5B**). Live imaging revealed that both tBB*ζ* and 11D5-3BB*ζ* CAR T cells led to efficient target cell killing (**Figure 5C**) via stable immune synapse formation (**Figure 5D and 5E**). While non-signaling tΔ and 11D5-3Δ CAR T cells did not form synapses nor kill target cells, they formed significantly more loose contacts with target cells than tBB*ζ* and 11D5-3BB*ζ* CAR T cells, suggesting significant interactions that may lead to BCMA pathway activation (**Figure 5F**). The duration of these loose contacts was longer with 11D5-3Δ compared to tΔ CAR T cells (**Figure 5G**), potentially explaining why 11D5-3Δ CAR T cells, and not tΔ, activated BCMA pathway. These results suggest that significant CAR T cell - tumor cell interactions can take place in the absence of killing which led us to hypothesize that those interactions could lead to a survival advantage of residual BCMA positive tumor cells.

**Figure 5.**
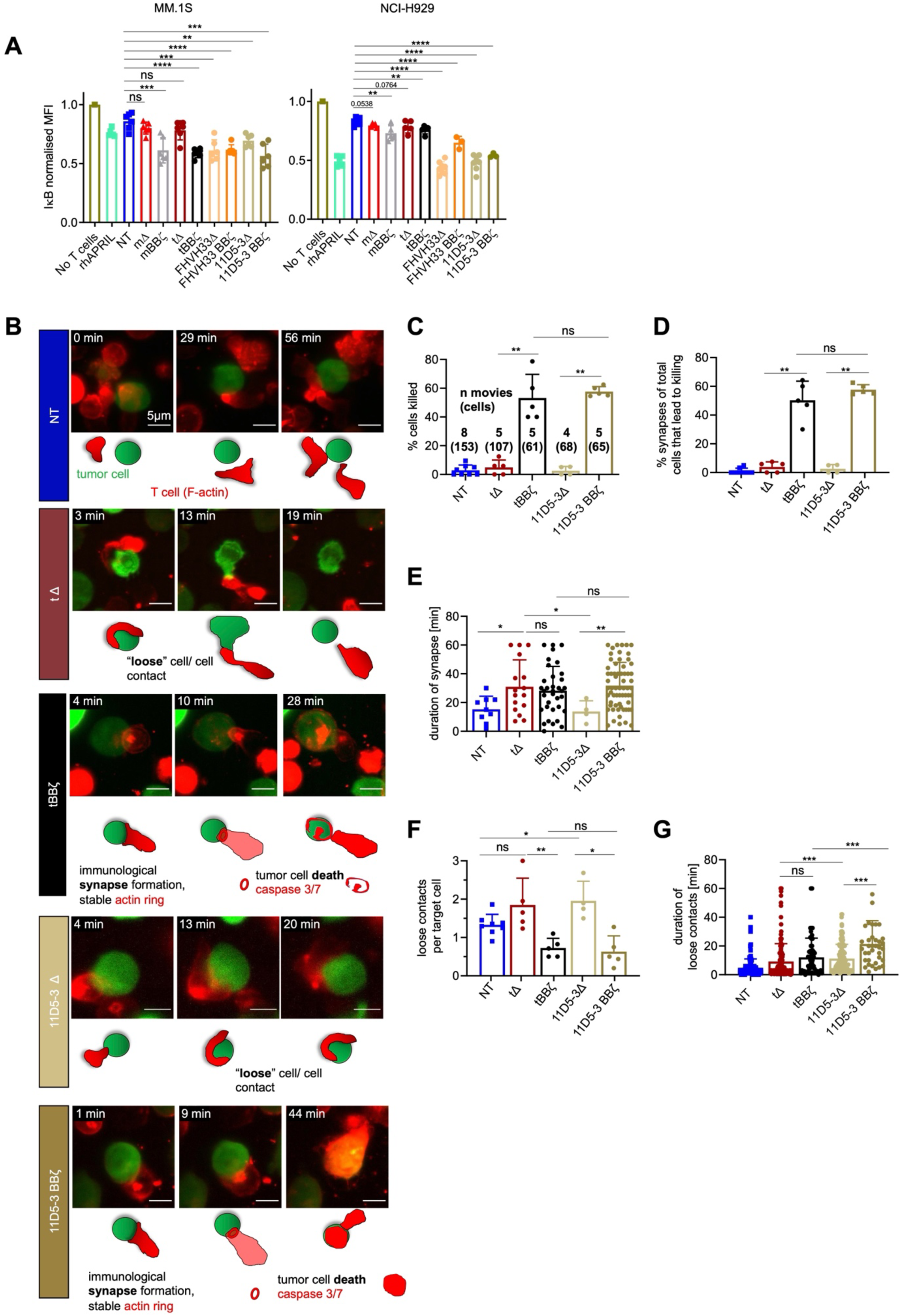
NFkB pathway activation in MM cells upon interaction with CAR T cells. (**A**) Normalized intracellular IκBα MFI in MM cells co-cultured with CAR T cells for 30 min. MM.1S cells (left, E:T 1:1), NCI-H929 cells (right, E:T 10:1). n=3-6 donors, mean±SD, unpaired t-test. (**B-G**) Light sheet live imaging of immunological synapse formation, target cell killing and loose T cell – tumor cell interactions. (**B**) Representative stills from light sheet time-lapse movies. NT (dark blue), tΔ (dark red), tBB*ζ* (black), 11D5-3Δ (beige), and 11D5-3BB*ζ* (dark beige) CAR T cells with MM.1S-GFP-FFLuc (green) target cells. T cell F-actin (SPY-actin live dye (red)), and cell death (biotracker caspase 3/7 stain, red). One image per minute for 60 min. Scale bar 5µm. (**C**) Percent target cell killing per 60 min movie, mean±SD. Numbers of movies and numbers of single cells (in parentheses) analyzed from n=3 independent donors and experiments. (**D**) Percentage of observed immunological synapses between T cells and target cells (appearance of an F-actin ring stable for > 2 minutes leading to target cell death measured by biotracker caspase 3/7 red signal), mean±SD. (**E**) Contact time between T cell and target cell at each synapse in minutes, mean±SD. (**F**) Quantification of loose contacts per target cell, mean±SD. (**G**) Contact time between T cell and target cell in minutes for loose contacts, mean±SD. (**A, C-G**) Definition of significance levels: ns=not significant, *p<0.05, **p<0.01, ***p<0.001, **** p<0.0001.

### Non-functional BCMA-directed CAR T cells do not promote tumor progression in vivo

Soluble APRIL is well known to promote tumor progression by activation of BCMA signaling in MM cells.^23-28^ Thus, we next investigated if APRIL-, FHVH33-or 11D5-3 ΔCAR T cells mediated MM cell proliferation. We showed that CAR T cells did not induce proliferation of MM.1S or NCI-H929 cells in vitro (**Figures 6A and 6B)**.

**Figure 6.**
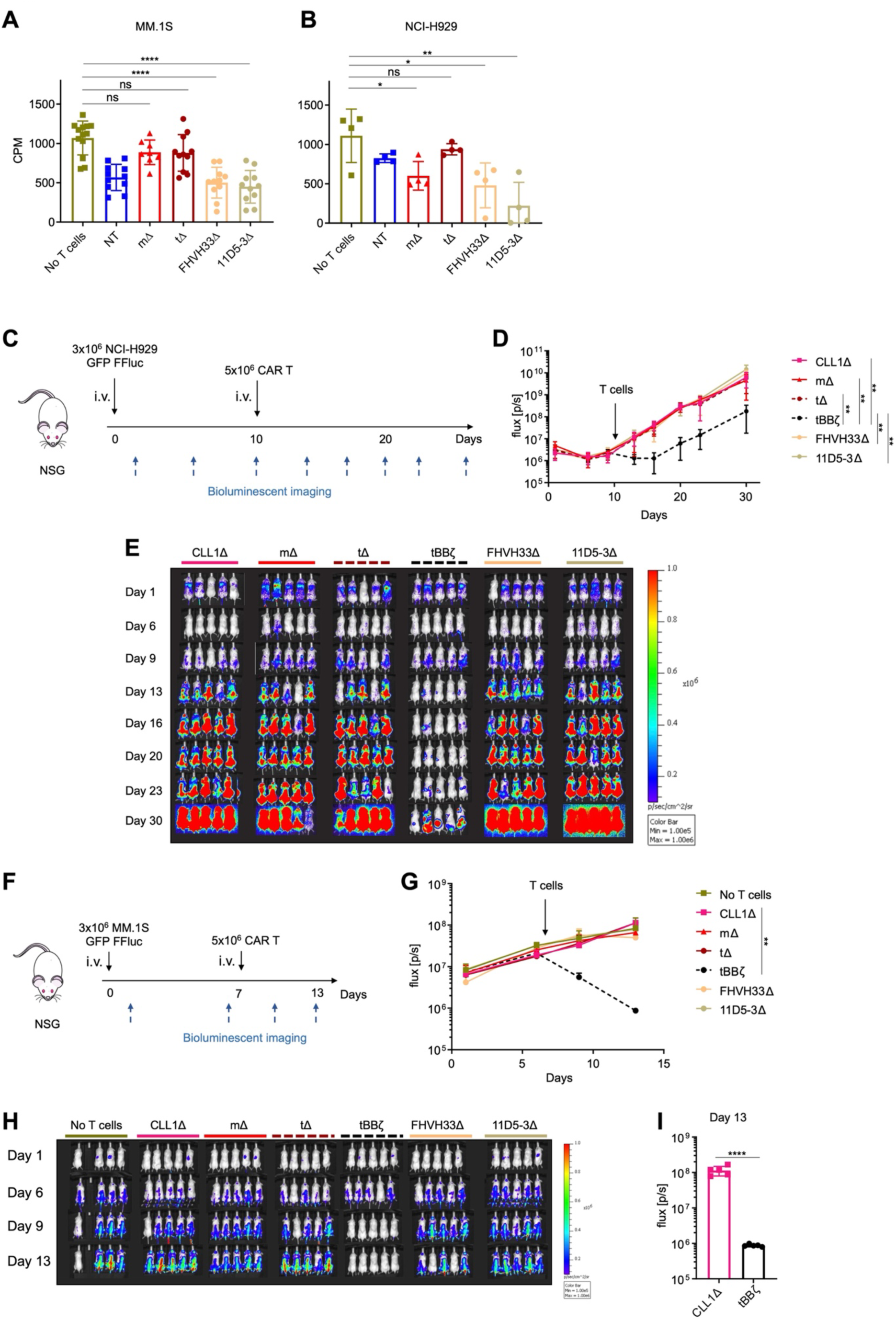
Non-functional BCMA-directed CAR T cells do not promote tumor growth. (**A, B**) H3 thymidine incorporation assay upon CAR T cell co-culture with (**A**) MM.1S at E:T ratio 1:1 or (**B**) NCI-H929 at E:T ratio 5:1. Background T cell H3 thymidine incorporation was subtracted. n=4-11 donors, colored symbols depict mean of technical triplicates, bars mean±SD, unpaired t-test. (**C**) Scheme of the NCI-H929 mouse xenograft model. 1 representative experiment of 2. n=5 mice per group per experiment. (**D**) BLI summary and quantification of total flux [p/s], mean±SD, unpaired Mann-Whitney-U test on AUC. (**E**) Tumor growth measured by BLI, individual mouse pictures (color scale min 1×10^5^, max 1×10^6^ p/sec/cm^2^/sr). Dorsal view identifies tumor growth in the spine and skull. (**F**) Scheme of the MM.1S mouse xenograft model. 1 representative experiment of 2. n=4-5 mice per group per experiment. (**G**) BLI summary and quantification of total flux [p/s], mean±SD, unpaired Mann-Whitney-U test on AUC. (**H**) Tumor growth measured by BLI, individual mouse pictures (color scale min 1×10^5^, max 1×10^6^ p/sec/cm^2^/sr). Dorsal view identifies tumor growth in the spine and skull. (**I**) Bar graph shows comparison of BLI signal intensity at day 13 between CLL1Δ and tBB*ζ* CAR T cell treated mice. (**A, B, D, G, I**) Definition of significance levels: ns=not significant, *p<0.05, **p<0.01, ***p<0.001, **** p<0.0001.

Similarly, in vivo in two different mouse xenograft models, we found no tumor promoting capacity of APRIL-, FHVH33- or 11D5-3 ΔCAR T cells, neither in mice systemically engrafted with NCI-H929 cells (**Figures 6C-E**) nor in mice systemically engrafted with MM.1S cells (**Figures 6F-H**). We confirmed significant anti-tumor activity with tBB*ζ* CAR T cells in both models (**Figures 6C-I**). Thus, despite significant T cell - tumor cell interactions that activated BCMA signaling, non-functional APRIL-, FHVH33- or 11D5-3 CAR T cells did not promote tumor growth in vivo.

## Discussion

We show that a trimeric configuration of truncated APRIL can be exploited as efficient binding moiety in CARs leading to dual antigen recognition on MM. Polyfunctionality and immune synapse formation were superior in CAR T cells with our trimer-compared to monomer-derived APRIL. The tBB*ζ* construct was most potent in vivo in two different MM mouse xenograft models and led to fast anti-tumor responses, but tumors recurred at later time-points. We identified BCMA downmodulation as an escape mechanism for APRIL-, scFv- and FHVH-CAR T cells that was attributed in part to trogocytosis and mostly to BCMA internalization. Despite activation of survival pathways in residual tumor cells and significant T cell – tumor cell interactions via loose contacts, we demonstrate that APRIL-, scFv-, and FHVH-CAR T cells cannot promote tumor cell proliferation or tumor growth in vitro and in vivo.

Truncated APRIL for the generation of natural ligand-based CARs targeting MM has previously attracted attention due to APRIL’s natural high affinity for BCMA and TACI expressed on MM, allowing for dual antigen targeting.^7 13^ In addition, both BCMA and TACI provide survival and proliferation signals to MM cells and are expressed on putative MM stem or progenitor cells.^6 25 28^ While the APRIL monomer based CD28-OX40*ζ* CAR showed promising preclinical activity,^7^ the subsequent clinical trial was terminated early due to insufficient efficacy in patients.^29^ A trimeric APRIL configuration as CAR binding moiety linked to a CD8*α* hinge and transmembrane and a BB*ζ* endodomain (TriPRIL) resulted in better polyfunctionality compared to a monomeric APRIL BB*ζ* CAR.^13^ Both groups demonstrated in their preclinical models that APRIL CARs have the capacity to overcome BCMA negative antigen escape, as mice bearing BCMA negative TACI positive myeloma xenografts could be controlled with TriPRIL CAR T cell therapy in vivo. A clinical trial assessing safety and efficacy of TriPRIL CAR T cells has recently started.^30^ Here, we investigated alternative APRIL CAR configurations and found that a histidine residue in position 115 of APRIL is potentially favoring the proper trimer folding and stabilization of the trimeric binding module. Linker length is also critical for the stability of the fold. We show that immune synapse formation is enhanced in tBB*ζ* compared to mBB*ζ* CAR T cells, correlating with polyfunctionality by intracellular cytokine staining. In vivo in the mouse xenograft models, tBB*ζ* CAR T cells produced significantly better anti-tumor control than mBB*ζ* CAR T cells, even though tumors recurred in the follow up. Overall, APRIL is an interesting natural dual antigen binder that can be used as a targeting moiety in CARs. Additional optimization is most likely required to deploy its full therapeutic potential.

CAR T cell therapy in MM has so far mostly focused on targeting BCMA as a single antigen. While fast and deep responses have been consistently reported in several clinical trials, most patients do not experience long-term sustained remissions.^10 22 31-35^ Mechanisms of resistance to BCMA CAR T cell therapy have not been entirely elucidated yet.^36^ Inspired from the CD19 CAR T cell experience, tumor escape by target antigen loss was already investigated. Indeed, rapid BCMA downmodulation on residual plasma cells has been reported in several clinical trials of BCMA-CAR T cell therapy, but it is not known if the BCMA antigen density on persisting myeloma cells is below the antigen sensitivity of the respective CAR T cells.^22 31^ In most patients, malignant plasma cells will re-express BCMA to higher levels on their cell surface upon disease progression, suggesting that BCMA downmodulation is a transient effect. Only two cases of definitive loss by genetic deletion have been described to date,^9 11 12^ probably because BCMA provides crucial signals for maintaining and expanding the malignant myeloma cell population.^28^ Our data now indicate that BCMA downmodulation consistently occurs in malignant plasma cells upon CAR T cell exposure, and we identified two mechanisms that contribute to this effect. First, CAR T cells can extract BCMA from their target cells and present it on the T cell surface. This phenomenon is known as trogocytosis and can potentially lead to CAR T cell exhaustion and fratricide, and thus limit CAR T cell function and persistence in vivo.^20^ Second, plasma cells internalize BCMA upon CAR T cell exposure, leading to a rapid reduction of BCMA cell surface levels. Similar internalization has been observed upon exposure to an antibody-drug conjugate targeting BCMA.^32^ Interestingly, several clinical trials indicate that a significant proportion of patients experienced disease progression despite peripheral blood CAR T cell persistence.^9 22^ Thus, other mechanisms of therapy resistance may include the limited fitness or exhaustion of persisting CAR T cells, limited trafficking of CAR T cells to the bone marrow, local immune suppression of CAR T cells in the tumor microenvironment, or BCMA pathway stimulation by continuous but insufficient CAR T cell – tumor cell interactions. In preclinical models, we investigated whether persisting non-functional mAPRIL-, tAPRIL-, 11D5-3, and FHVH33-CAR T cells could stimulate the BCMA pathway and through this interaction promote tumor growth. Even though NFkB signaling was activated upon loose T cell - tumor cell contacts with all CARs investigated except with mΔ and tΔ, this stimulation did not translate into measurable MM cell proliferation in vitro or faster tumor progression in vivo in two mouse xenograft models. Thus, these findings are encouraging and suggest that BCMA targeted CAR T cells are not actively involved in tumor progression even if they persist in vivo in a non-functional or exhausted state. Even though soluble APRIL is a strong survival factor involved in MM progression and high levels in patients are associated with poor prognosis,^26 37^ truncated APRIL presented on non-functional CAR T cells did not promote tumor progression. A potential explanation could be that we used a truncated sequence devoid of the heparan sulfate proteoglycan binding (HSPG) site that is required to engage CD138 on plasma cells and critical for mediating the pro-proliferative and survival effects mediated by soluble APRIL.^38^

Due to the importance of BCMA in MM cell biology, BCMA remains a very attractive target for CAR T cell therapy. Strategies to upregulate BCMA levels on target cells before and early during the course of CAR T cell therapy are investigated clinically by the use of GSI.^21 39^ In our study however, when adding GSI simultaneously with CAR T cells to the co-culture, we nevertheless found BCMA downmodulation to a similar extent as in the absence of GSI. A pretreatment phase with GSI seems to be critical to achieve the desired effect on increasing BCMA levels on target cells.^21^ Another avenue for improvement is the use of fully human binding sequences to reduce immunogenicity of CAR T cells, as we do with APRIL CAR T cells. In clinical trials using murine scFvs in CARs such as the 11D5-3 scFv, anti-drug antibodies (ADAs) were detected in a significant proportion of patients,^9 33^ and the presence of ADAs was a risk factor for relapse or progression after CAR T cell therapy.^33^ Further, efficacy may be increased with dual antigen targeting CARs that combine BCMA targeting with a second antigen that is involved in myeloma-genesis but through a different pathway. Combinatorial targets include explorations of GPRC5D, SLAMF7, CD38, CD19, TACI and BAFF-R.^40-47^ Other future directions include combinatorial targeting with CAR T cells and microenvironment modulation or the exploration of enhanced manufacturing modalities that favor the long-term function and persistence of adoptively transferred T cells.

In summary, we show that trimeric APRIL CAR T cells can efficiently target both BCMA and TACI on MM, and that our tBB*ζ* construct was most potent. We reveal BCMA downmodulation on tumor cells as an important mechanism of immune evasion upon CAR T cell contact mediated by trogocytosis and BCMA internalization, independent of the binding moiety used. Importantly, when non-functional mAPRIL, tAPRIL, 11D5-3 or FHVH33 CAR T cells persisted in vivo, they were not able to promote tumor proliferation and growth in vivo. Our results shed light on the mechanisms underlying CAR T cell treatment failure targeting BCMA in MM and may help to devise more efficient therapeutic strategies in the future.

## Acknowledgements

We are very grateful to Maude Varrin for technical assistance and lab management, to Dr. Rosa Paolicelli, University of Lausanne, for granting us access to her light sheet microscope for live imaging, and to Francisco Sala de Oyanguren for expert technical assistance with the Image Stream analysis. We thank all Arber lab members for helpful and stimulating discussions and critical feedback on the manuscript.

## Author contributions

N.C. designed research, performed experiments, analyzed and interpreted results, and wrote parts of the manuscript. B.W. designed research, performed microscopy studies, and wrote parts of the manuscript. G.C. performed statistical analysis of polyfunctionality assessment. D.G. supervised statistical analysis of polyfunctionality assessment. V.Z. designed research, performed molecular modeling, analyzed and interpreted results and wrote parts of the manuscript. C.A. designed research and supervised the entire study, analyzed and interpreted results and wrote the manuscript. All authors reviewed and approved the final version of the manuscript.

## Data availability

The data that support the findings of this study are available from the corresponding author upon reasonable request.

