## supplemental material and figures for "Rapid BCMA downmodulation on myeloma cells upon CAR T cell contact is mediated by trogocytosis and BCMA internalization"

**Running title:** Mechanisms of CAR T cell failure in myeloma

### Materials and Methods

**Cell lines.** MM.1S (multiple myeloma) and 293T (human embryonic kidney) cells were purchased from the American Type Culture Collection (ATCC). K562 (chronic myeloid leukemia) and NCI-H929 (multiple myeloma) were purchased from the German Cell Culture Collection (DSMZ). KG-1a (acute myeloid leukemia) cells were a gift from Dr. Stephen Gottschalk, Center for Cell and Gene Therapy, Baylor College of Medicine, Houston, USA. K562 and MM.1S were maintained in RPMI 1640 (Gibco, # 61870010) with 10% Fetal Bovine Serum (FBS) (Gibco or Biowest). KG-1a were maintained in RPMI 1640 with 20% FBS. NCI-H929 were maintained in RPMI 1640 with 20% FBS, 1% sodium pyruvate (Gibco) and 50µM β2-mercaptoethanol (Gibco). 293T cells were maintained in Dulbecco's modified Eagle medium (DMEM, Gibco, #21885025) supplemented with 10% FBS. All cell culture media also contained 1% penicillin-streptomycin (Gibco), and 1% L-glutamine (Bioconcept) (**Supplementary Tables 1 and 2**). The PG-13 retroviral producer cell line for the generation retroviral particles encoding Green Fluorescent Protein (GFP) and firefly luciferase (FFLuc) (GFP-FFLuc) was kindly provided by Dr. Stephen Gottschalk, Baylor College of Medicine. MM.1S and NCI-H929 cells were retrovirally transduced with GFP-FFLuc and FACS sorted for >98% purity. K562 cells were engineered to express TACI ($K562_{TACI+}$), BCMA ($K562_{BCMA+}$) or BCMA and TACI ($K562_{BCMA+TACI+}$) by retroviral transduction, and FACS sorted for >98% purity. Cell lines were authenticated in January 2022 by Microsynth.

**APRIL CAR modelling.** The 3D structure of the soluble human APRIL monomer, trimer or APRIL-based CARs in complex with BCMA and TACI were obtained by homology modelling using the experimental structure of mouse APRIL bound to human TACI or BCMA (PDB ID 1xu1 and 1xu2, respectively[1]). The sequence alignment between mouse APRIL, human APRIL and the CAR proteins were obtained using the MUSCLE program.[2] Based on this sequence alignment, 2000 structural models were generated using the MODELLER program[3] for each APRIL/ligand combination, and ranked according to the DOPE energy score.[4] The top-ranked models according to DOPE were retained as the final model. Molecular visualization and analysis were performed using the UCSF Chimera software.[5]

The solvent accessible surface area (SASA) of the CAR-trimer:BCMA complex, the isolated CAR-trimer and the BCMA protein were calculated using the analytical procedure of the CHARMM v43 program, using the CHARMM v27 forcefield (https://www.charmm.org). The SASA of the CAR-trimer:BCMA complex was obtained from the cartesian coordinates of the corresponding homology model, while those CAR-trimer and the BCMA protein were obtained from the same starting coordinates, after removing the BCMA and CAR proteins, respectively. The same procedure was used to calculated the SASA of the CAR-monomer:BCMA complex, of the isolated CAR-monomer and of the corresponding isolated BCMA protein, and for the human model of the APRIL-trimer:BCMA complex, isolated APRIL-trimer and the corresponding isolated BCMA protein.

**Generation of retroviral vectors.** Plasmids encoding for human truncated APRIL (115-250 a.a, Uniprot O75888, excluding the heparan sulfate proteoglycan binding sites of APRIL[6 7]), full-length BCMA (Uniprot Q02223), full-length TACI (Uniprot O14836), 11D5-3 single chain variable fragment (scFv[8]) and the human heavy-chain-only FHVH33[9] were codon-optimized and synthesized by GeneArt (Thermo Fisher Scientific). Retroviral constructs encoding APRIL CARs, 11D5-3 and FHVH33 CARs, BCMA, TACI and BCMA-Emerald were generated using the In-Fusion HD Cloning Kit (Takara, #638933) according to manufacturer's instructions. Sequences of interest were amplified by high-fidelity PCR (CloneAmp™ HiFi PCR Premix, #639298). p-QGXIP retroviral backbone (kindly provided by Dr. Melita Irving, Department of Oncology UNIL CHUV, University of Lausanne, Switzerland) was linearized by EcoRI restriction enzyme digestion for the cloning of BCMA. p-SFG retroviral backbone was linearized by NcoI and NotI restriction enzyme digestion for TACI cloning. p-SFG retroviral backbone containing an IRES-ΔCD271 reporter gene (**Figure 1E**) was linearized by PmeI and XhoI restriction enzyme digestion for CAR cloning (**Supplementary Table 3**). For CAR cloning, 11D5-3, FHVH33, APRIL, CD8 hinge and transmembrane domain[10], 41BB[10], CD28[10] and CD3ζ[10], sequences were amplified using overlapping primers designed with In-Fusion Cloning Primer Design Tool. PCR fragments were gel purified from an agarose gel using the QIAquick Gel Extraction Kit (Qiagen, #28706X4). Fragments of interest were assembled using the In-Fusion enzyme mix with the linearized backbone to generate the constructs of interest and transformed into stellar competent cells (Takara, #636763). Plasmid DNA was purified from minipreps with QIAprep spin Miniprep Kit (Promega), and constructs were verified by sequencing (Microsynth).

**Generation of retroviral supernatant.** Retroviral supernatant was produced by transient transfection of 293T cells as previously described.[11] In brief, 293T cells at 50% confluency were co-transfected with 1) the RDF plasmid encoding the RD114 envelope or a plasmid encoding the VSVg envelope, 2) the Peg-Pam plasmid encoding MoMLV gag-pol or a plasmid encoding gag-pol, and 3) the SFG retroviral plasmid of interest or QGXIP retroviral plasmid (with LTRs and packaging signals), using GeneJuice transfection reagent (Merck, #70967-3) according to manufacturer's instructions. Retroviral supernatants were harvested after 48 and 72 hours of culture, filtered with 0.45μM filter (Filtropur S, Sarsdedt #83.1826), snap-frozen on a dry ice/ 100% ethanol mixture, and then stored at -80°C until use, or used as fresh supernatant.

**Peripheral blood mononuclear cells from healthy human donors.** Buffy coats from de-identified healthy human volunteer blood donors were obtained from the Center of Interregional Blood Transfusion SRK Bern (Bern, Switzerland).

**Generation of CAR T cells and transgenic cell lines.** Peripheral blood mononuclear cells (PBMCs) were isolated from buffy coats by density gradient centrifugation (Lymphoprep, StemCell #07851). PBMCs were activated on plates coated with anti-CD3 (1mg/ml, Biolegend, #317347, clone: OKT3) and anti-CD28 (1mg/ml, Biolegend, #302934, clone: CD28.2) antibodies in T cell media (RPMI containing 10% FBS, 2mM L-Glutamine, 1% Penicilin-Streptomycin) with IL-15 and IL-7 (Miltenyi Biotec, 10ng/ml each,

#130-095-362 and #130-095-765, respectively) (**Supplementary Table 1**). The day before transduction, a non-tissue culture treated 24-well plate (Grener Bio one, #662102) was coated with retronectin (Takara Bio, #T100B) in PBS (7µg/ml, 1ml per well), and incubated overnight at 4°C. Three days after activation, retronectin was removed and the plate was blocked with T cell medium during 15 min at 37°C. Then, media was removed, and retroviral supernatant was centrifuged onto the retronectin coated plate at 2000g for 1hour at 32°C. Retroviral supernatant was gently removed and activated T cells at $0.15x10^6$ cells/ml was added, and centrifuged at 1000g for 10 min at 21°C. Cells were then incubated at 37°C/ 5% $CO_2$ for 3 days. For the generation of transgenic cell lines, cells were counted the day of transduction (0.15x10^6 cells/ml) and performed as described for T cells. After 48 to 72 hours of transduction, T cells were harvested and further expanded in T cell media containing IL-7 and IL-15. In selected experiments, CAR T cells were positively selected with a CD271 selection kit (EasySep, #17849) to enrich for transduced T cells. Transgenic cell lines were then expanded in the appropriate media, checked by flow cytometry for BCMA and/or TACI, or GFP expression, and FACS sorted.

**Sequential co-culture assay.** K562, $K562_{BCMA+}$, $K562_{TACI+}$, $K562_{BCMA+TACI+}$, KG-1a, NCI-H929 and MM.1S, expressing or not GFP-FFLuc, were co-cultured with T cells in five replicate wells at an effector to target (E:T) ratio of 1:1 (200'000 cells each in a 48-well plate). After three to four days, residual tumor cells and T cells were quantified by FACS combining antibody staining and absolute counting beads (CountBright Beads, Life Technologies #C36950). Fresh tumor cells were added back to replicate wells when

>85% of tumor cells were killed at the analysis timepoint. Otherwise, the killing was considered incomplete and T cells were not rechallenged with fresh tumor cells.

**Immunophenotyping and FACS sorting.** Cell surface stainings were performed using antibodies (**Supplementary Table 5**) during 15min at 4°C, or RT during 20min for CCR7 staining. Other flow cytometry related reagents are listed in **Supplementary Table 6**. Cell viability was assessed with DAPI (Biolegend) or Zombie UV (Biolegend). CAR cell surface was performed by APRIL staining using a primary biotinylated antibody against APRIL (Biolegend) followed by Streptavidin-APC (Biolegend or Biorad).

To evaluate CAR T cell polyfunctionality, protein transport in T cells was blocked with golgiplug (BD, #555020, diluted 10x in PBS, 5µl/ml) and golgistop (BD, #554724, diluted 10x in PBS, 5µl/ml), and CAR T cells were stimulated with different tumor target cell lines (K562, $K562_{BCMA+}$, $K562_{TACI+}$ or $K562_{BCMA+TACI+}$) in a 1:1 ratio. The anti-CD107a antibody was added at the beginning of the co-culture (10µl/ml). After 8 hours of stimulation, cells were collected and cell surface markers were stained. Then, cells were fixed and permeabilized (Cytofix/Cytoperm, BD, #554714) during 20min at 4°C in dark. Cells were washed once with FACS buffer (Phosphate buffered saline (PBS) containing 2% FBS, and 0.1% Saponin (Sigma, #S7900-25G)), and intracellular staining was performed during 30 min at 4°C in dark. Cells were washed twice with FACS buffer 0.1% Saponin, resuspended in FACS buffer and analyzed. Data was visualized with SPICE software (version 6.0) after extracting polyfunctional cell populations from Flowjo, and converting to a file format using Pestle software (version 2.0).

To assess NFκB pathway activation, MM cells (MM.1S or NCI-H929) were rested overnight in pure RPMI 1640. The next day, cells were exposed to recombinant human APRIL (400ng/ml, Peprotech, #310-10C), CAR T cells, or media alone for indicated times. Cells were then fixed immediately with Cytofix (BD, #554655) during 10 min at room temperature (RT) followed by permeabilization with Perm Buffer III (Phosflow, BD, #558050) on ice for 30 min. Staining for intracellular IκBα was performed at RT for 60 min with anti-IkBα-PE antibody (eBioscience, #12-9036-42, clone: MFRDTRK).

Sample acquisition was performed on a BD FACS LSR II, SORP or Fortessa, using DIVA software (version 8.0.1). Data analysis was done using FlowJo software (FlowJo V10 or higher). FACS sorting to purify transduced cell lines was performed by the flow cytometry facility (University of Lausanne, Biopole 3, Epalinges, Switzerland) using a FACSARIA IIu instrument and DIVA software. Cell lines engineered with GFP-FFluc, were sorted based on GFP+ gating, cell lines engineered with BCMA were sorted after staining with anti-BCMA-PE, cell lines engineered with TACI were sorted after staining with anti-TACI-PE.

To analyze cell surface co-localization of anti-BCMA PE antibody and Emerald, MM.1S-BCMA-Emerald cells were first incubated during 1h with T cells. After surface staining with anti-BCMA PE antibody (Biolegend, #357504, clone 19F2), cells were fixed with Cytofix (BD, #554655) and analyzed on an Amnis ImageStreamX Mark II instrument. Samples were run in a 2 camera, 12 channel ImageStreamX multispectral imaging flow cytometer (Luminex Corporation) at low speed and highest magnification (60X). Instrument setup and performance tracking was performed daily using the Amnis® SpeedBead® Kit (Luminex Corporation) for verifying optimal instrument performance. Cells were excited using a 488 nm laser (40mW) and a 561nm yellow laser (200mW).

Only events with a brightfield area greater than 50 µm² (to exclude cell debris) and non-saturating pixels were collected (as described in[12]). Data were acquired for 30,000 events/sample. Experimental samples contained images and data for Brightfield (Channels 1 and 9), Emerald (Channel 2), BMCA-PE (Channel 3) and SSC (Channel 12). Single color controls for each fluorochrome were acquired to generate the compensation matrix that was applied to all the experimental files prior to analysis using IDEAS (Image Data Exploration and Analysis Software) 6.2 software (Luminex Corporation). After compensation, similarity analysis was done on in-focus single cell images. In-focus single cells were classified based on their brightfield area, high brightfield aspect ratio (width to height ratio) and low SSC signal. Colocalization of fluorescent probes was measured in the population of double positive cells for Emerald and PE using the Bright Detail Similarity feature. This feature quantifies the degree of correlation between any two channels images on a pixel-by-pixel and cell-by-cell basis. It is derived from the Pearson's correlation coefficient (ρ), which is based on a linear regression analysis of pairs of values taken from different data sources. The data pairs are the pixel intensities at the same location (x,y) in each image of the Emerald and PE channels respectively. The score is negative in sign for images that are opposites, close to zero for uncorrelated images and positive for similar images. High values (≥1) indicate that the fluorescence of Emerald and PE appear in the same location (same pixel coordinates).

$$\text{Similarity} = \ln\left(\frac{1+\rho}{1-\rho}\right) \qquad \rho = \frac{\sum_i (x_i - X)(y_i - Y)}{\sqrt{\sum_i (x_i - X)^2 \sum_j (y_j - Y)^2}}$$

The BDS score is derived from the Pearson's correlation coefficient (ρ). X and Y are the corresponding mean intensity values; xi and yi are the per pixel intensity values of the two images.[12 13]

**ELISA and MesoScale Discovery Assay (MSD).** Soluble BCMA (sBCMA) was detected in culture supernatants using a commercial ELISA kit (Human BCMA/TNFRSF17 DuoSet ELISA, R&D Systems, #DY193). Plate measurement was done on a SpectraMax M3 spectrophotometer, and data was acquired on SoftMax Pro software (version 6.2.2). Data were analyzed in Microsoft Excel (2016). The blank value was subtracted from all values acquired, and the standard curve was used to calculate sBCMA concentrations in ng/ml in each sample. For tonic signaling experiments, CAR T cells were cultured for 24h in media without cytokines, and culture supernatants were harvested. IL-2 concentrations in supernatants were analyzed in duplicates with the MSD MesoScale assay and data analyzed in Microsoft Excel (2016).

**Proliferation assay.** MM cell proliferation was assayed by 3H thymidine (Perkin-Elmer) incorporation. MM cells ($2x10^4$ per well) were cultured alone or in presence of T cells ($2x10^4$ per well) in triplicate wells on a 96-well round bottom plate at 37°C and 5% CO2 during 4 days. 3H Thymidine (0.5µCi/well) was added during the last 16h of culture. Cells were then harvested on MultiScreen Filter plates (Merck Millipore, # MAHFC1H60), and the counts per minute were measured on a TOPcount NXT (Serie number: B9912V, Perkin Elmer) instrument. The data were acquired on the TopCount NXT software version

1.06. Data analysis was done in Microsoft Excel (2016), and the background of T cell proliferation was subtracted to wells with MM cells and T cells in co-culture.

**Mouse xenograft experiments.** All animal studies were conducted in accordance with a protocol approved by the Veterinary Authority of the Swiss Canton of Vaud and performed in accordance with Swiss ethical guidelines. NOD-SCID-$\gamma c^{-/-}$ (NSG) mice were bred and maintained at the animal facility of the University of Lausanne, site of Epalinges. Animals were 7-11 weeks old at the start of the experiments, both males and females were used, and experimental groups were randomized based on animal weight at start of experiment.

For the NCI-H929 model, NSG mice were injected with $3x10^6$ NCI-H929-GFP-FFluc cells intravenously (i.v.) in the tail vein on day 0. Two adoptive T cell transfers were performed with $5x10^6$ T cells i.v. on day 10 and $3x10^6$ T cells i.v. on day 20, or only one T cell transfer on day 10 with $5x10^6$ T cells since no additional benefit of a second infusion was seen (**Figure 2D**). Each experiment is outlined with an experimental scheme in the manuscript figures.

For the MM.1S model, NSG mice were injected with $3x10^6$ MM.1S-GFP-FFluc cells i.v. in the tail vein on day 0. One adoptive T cell transfer was performed with $5x10^6$ T cells i.v. on day 7.

Tumor growth was monitored twice a week or weekly by bioluminescent imaging (BLI) on a Xenogen IVIS Lumina II instrument using Living Image software (version 4.7.3, 64bit). Data analysis was performed on the same software. Survival was determined according to humane endpoints as defined in the score sheet of the approved animal protocol.

**Immunofluorescence staining of fixed cells.** 12 mm glass coverslips (Thermofisher, #CB00120RA120MNT0) were coated overnight with 0.1mg/ml of Poly-D-Lysine (Thermofisher, #A3890401). After three washes with 1% PBS MM.1S GFP-FFLuc cells were seeded onto the glass coverslips. After target cell adherence, usually around 30 min, an equal number of CAR T cells was added. After 30 min of incubation at 37°C, 5% CO2, cells were fixed for 8 min in 4% paraformaldehyde (ThermoScientific, #28909). Next, cells were washed in PBS with 0.05% Triton X-100 (AppliChem, #1388) (PBST). After blocking in 1% BSA in PBST for 1 h at room temperature, cells were incubated with primary antibodies at RT for 4h. After three washes in PBST for 5 min each, cells were incubated with secondary antibodies for 1h at RT, stained with 1 μg/ml Hoechst 33342 (Sigma Aldrich), washed three times 5 min in PBST, and mounted. Anti-F-actin 546 (Chrometra) was added 1:1000 with the secondary antibody. Primary antibodies were 1:1000 mouse anti-GFP (Thermo Scientific, # A-11120) and 1:200 mouse anti-alpha Tubulin (SantaCruz, DM1 α, # sc 32293). Secondary antibody was Alexa Fluor 488–coupled anti–mouse (Life Technologies, #A-11029), used at 1:1000.

**Confocal imaging of fixed cells**. Image stacks with a distance of 0.4 μm between optical slices were acquired with a 60x, NA 1.42 plan apochromat oil objective at 526x526 pixel minimal resolution on an Oympus FluoView 3000 confocal microscope equipped with a charge-coupled device camera (Olympus DP80) at the Cellular Imaging Facility of the University of Lausanne. Images were then processed and analyzed in ImageJ (National Institutes of Health). Relevant optical slices are shown as maximum intensity projections.

**Light Sheet Live microscopy.**

*Imaging set up.* Live cell imaging was performed on the InVi SPIM inverted light-sheet microscope (Bruker, Luxendo), access was kindly granted by Prof. Rosa Paolicelli, University of Lausanne. The imaging chamber acts as an incubator with environmental control and it has a reservoir for immersion medium, which is filled with water so that both objective lenses and the bottom of the sample holder are below the water surface. The microscope is prepared 12 hours before imaging to let the system reach equilibrium conditions to avoid stage shifts during acquisition. Invi dishes (Ibidi via Bruker, #80-0031-00-00) were coated overnight with 0.1mg/ml of Poly-D-Lysine (Thermofisher, #A3890401). After three washes with PBS, MM.1S-GFP-FFLuc target cells were seeded on light sheet dishes and incubated at 37°C, 5% CO2 until 50% confluency of a single cell layer on the light sheet dish was reached (usually about 20 min). F-actin was labelled in CAR T cells and non-transduced (NT) cells using SPY555-actin (Spirochrome, #SC202) according to the manufacturer's instructions, minimum 45 min before imaging. Equal amounts of T cells were seeded on the target cells. To visualize cell death BioTracker NucView 530 Red Caspase 3/7 dye (PBS, # Merck, SCT105) was added to the light sheet dish (1uM final concentration). The light sheet dish was sealed with Parafilm and mounted to the sample holder.

*Imaging configuration and conditions.* The InVi SPIM is equipped with a Nikon CFI 10x/0.3NA water immersion lens for illumination and a Nikon CFI-75 25x/1.1NA water immersion lens for detection. For excitation of GFP and mCherry, 488 nm and 594 nm laser lines were used, respectively, while emission was selected using a 497–554 nm band pass filter and a 610 nm long pass filter, respectively. 3D image stacks were acquired with a light-sheet thickness of 4 mm, a final magnification of 62.5x, resulting in 104 nm pixel size. The InVi SPIM environmental control was set to 37˚C, 5% CO2 and 80% humidity. A series of optimization experiments, involving different laser powers, exposure times and z-step sizes yielded laser powers of 5 mW for 488 nm and 5 mW for 594 nm, 50 millisecond exposure time per frame and 4 μm z-spacing between frames to be optimal for imaging (96–120 hr) without photo-bleaching or photo-toxic effects on cells. Images were recorded as 50-100 2D planes with a framerate of 1 minute for a total of 1 hour in two channels equipped with filters for red (mCherry) and green (GFP) light.

*Image analysis*. Big Data Processor,[14] a Fiji plugin for lazy loading of big image data, was used to visualize the images in 2D slicing mode, crop stacks in x, y, z, and t, perform chromatic shift correction between channels and convert h5 files from the InVi SPIM into a Fiji compatible file format (tiff) for further analysis. Movies were analyzed manually and with a custom-made Fiji interactive macro to define and create regions of interest. Around 20 target cells were identified randomly per movie and condition. For those target cells the number of loose contacts with T cells and their duration was determined. Loose contacts were defined as lasting a minimum of 2 frames (2 minutes) and having the T cell clearly adopt to the shape of the target cell as a sign of loose surface molecule interaction. Immunological synapse formation between T cell and target cell was quantified as synapses per target cell. A synapse was defined as formation of an F-actin ring on the T cell/ target cell interface and a stabilized cell/ cell contact (no moving of the interface for at least 3 frames). Furthermore, the number of target cells dying per movie was determined using the biotracker red signal as a readout.

**Statistical analysis**. Descriptive statistics were used to summarize data. For continuous variables, normality distribution was analyzed, and comparisons were made by t-test. When normality testing failed, pair-wise comparison of conditions was performed using Mann-Whitney U-test. Area under the curve (AUC) comparisons were analyzed with unpaired t-test and Welch's correction (T cell expansion in vitro), or unpaired Mann-Whitney test (non-parametric t-test) (in vivo bioluminescent imaging). Sequential co-cultures and survival of mice were analyzed by Kaplan Meier method and significance assessed with log-rank (Mantel-Cox) test. Multiple comparisons (e.g. BCMA cell surface levels, analysis of sBCMA, T cell phenotypes) were performed by repeated Measures (RM) one-way ANOVA with Tukey's multiple comparison. Analyses were performed in GraphPad Prism version 8.1.2 or higher. Differences in T cell polyfunctionality was analyzed by dependent t-test for paired samples using Python scipy.stats package (version 1.5.4), where cells co-expressing three or more evaluated markers simultaneously were considered to be polyfunctional. P values $<0.05$ were considered significant, and the levels of significance are indicated in the figures and legends.

### Supplementary Figures and Legends

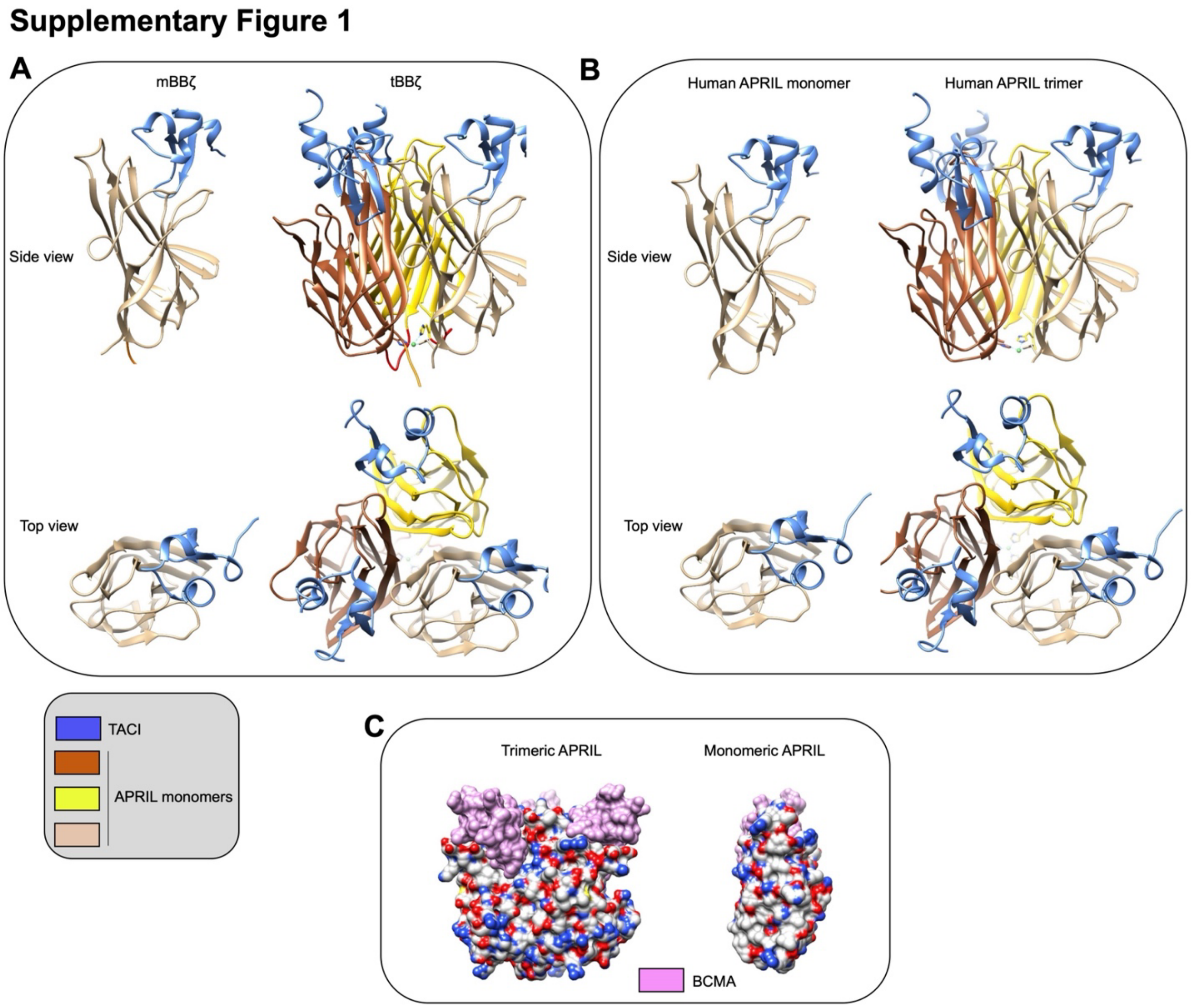


**Supplementary Figure 1. Molecular models of mAPRIL and tAPRIL CARs compared to soluble human APRIL interacting with TACI or BCMA.** (**A-B**) In silico modeling of monomeric and trimeric APRIL ligand binding domains used for mBBζ and tBBζ CAR constructs (**A**) interacting with TACI (blue), compared to natural APRIL protein (**B**). (**C**) Left: Surface of the APRIL trimer in complex with three copies of BCMA. BCMA surface is colored in pink, APRIL surface is colored as a function of the atom type (carbon in white, sulfur in yellow, nitrogen in blue and oxygen in red). Right: surface of a monomeric APRIL binding a single copy of BCMA. The solvent accessible surface area (SASA) of the APRIL trimer is composed by 45% of polar atoms (oxygen and nitrogen atoms), while the polar atoms constitute only 39% of the trimer SASA.

**Supplementary Text 1 (Figure 1 and Supplementary Figure 1).** We found that the average SASA that is buried upon binding of the CAR-trimer to BCMA is 1763 Å2, very close to the 1782 Å2 buried during the binding of the APRIL trimer to BCMA, but significantly larger than the 1521 Å2 buried upon the binding of the CAR-monomer to BCMA. Indeed, in the APRIL-trimer:BCMA and the CAR-trimer:BCMA complexes, although each of the three BCMA proteins bound to the APRIL trimer is mainly recognized by one monomer, several residues of a second monomer are contacting it, namely His203, Asp205, Arg206, Tyr208 and His241 (**Figure 1D**). These additional interactions are not present in the CAR-monomer:BCMA complex. Consequently, as could be expected, the CAR-trimer better recapitulates the physiological target recognition of the APRIL trimer than the CAR-monomer.

**Supplementary Text 2 (Figure 1 and Supplementary Figure 1).** It is well established that globular soluble proteins or protein complexes exhibit an increased concentration of polar residues at their surface (to help solubilize the protein), while their core is enriched in non-polar residues (to prevent non-favorable contacts with the polar water molecules). Indeed, we calculated that 45% of the SASA of the CAR-trimer, considering only the part corresponding to the APRIL trimer, is generated by the polar oxygen and nitrogen atoms. Of note, the removal of two of the monomers from the CAR-trimer to form the CAR monomer, exposes to the solvent a fraction of the surface of the remaining monomer that would normally be buried and solvent inaccessible in the trimer. Consequently, the fraction of the SASA of the CAR-monomer generated by the polar oxygen and nitrogen atoms, again considering only the part corresponding to the APRIL protein, drops to 39%. This could partly weaken the protein structure (**Supplementary Figure 1C**).

### Supplementary Figure 2

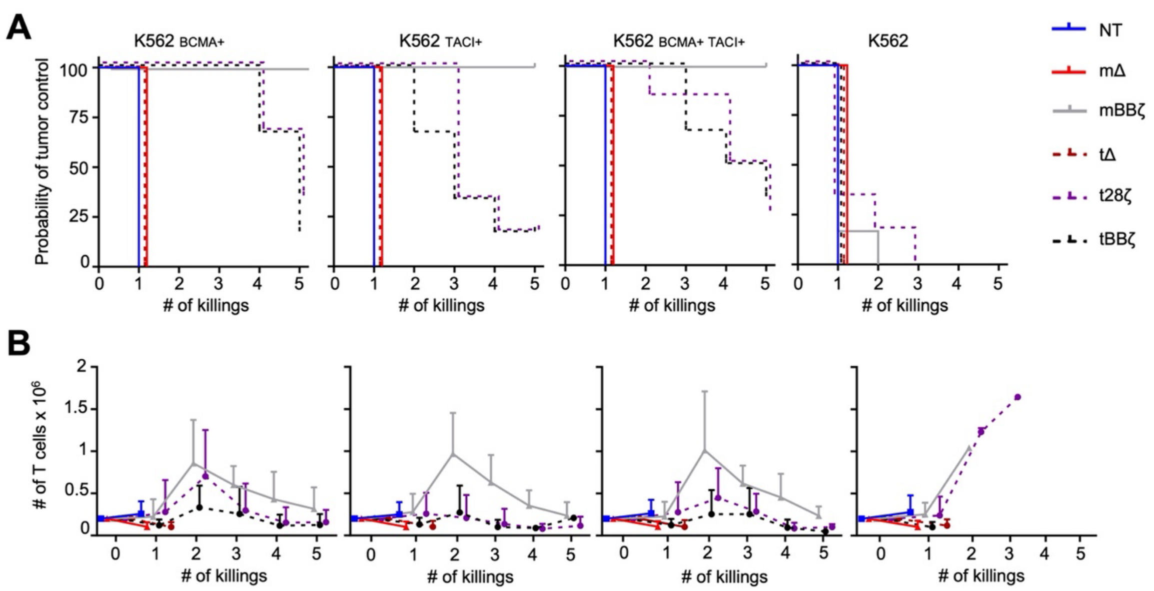


Log-rank test for probability of tumor control

| | K562 BCMA+ | K562 TACI+ | K562 BCMA+ TACI+ | K562 |
|---|---|---|---|---|
| mΔ vs mBBζ | *** | *** | *** | ns |
| mBBζ vs tBBζ | ** | ** | * | ns |
| t28ζ vs tBBζ | ns | ns | ns | ns |

T-test on area under the curve for T cell expansion

| | K562 BCMA+ | K562 TACI+ | K562 BCMA+ TACI+ | K562 |
|---|---|---|---|---|
| mBBζ vs tBBζ | * | ** | * | *** |
| t28ζ vs tBBζ | ns | ns | ns | ns |

**Supplementary Figure 2. In vitro anti-tumor function of APRIL CAR T cells against engineered K562 cells.** Sequential co-culture stress-test of APRIL CAR T cells with $K562_{BCMA+}$, $K562_{TACI+}$, $K562_{BCMA+TACI+}$ and K562 cells. (**A**) Kaplan Meier curves show the probability of tumor control, (**B**) the line graph depicts CAR T cell expansion. E:T 1:1, n=6 donors, mean±SD. Log-rank test to compare killing efficacy (left table), t-test on AUC to evaluate T cell expansion (right table). Significance levels: *p<0.05, **p<0.01, ***p<0.001.

### Supplementary Figure 3

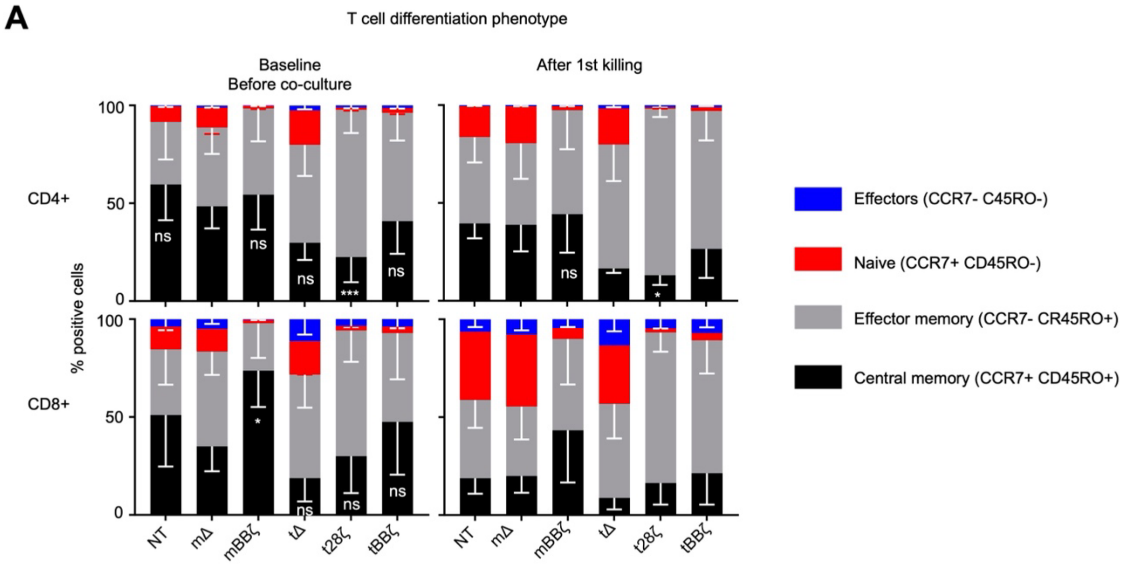


| | CD4 before co-culture | CD8 before co-culture | CD4 after 1st killing | CD8 after 1st killing |
|---|---|---|---|---|
| t28ζ vs tBBζ | ** | ns | ns | ns |

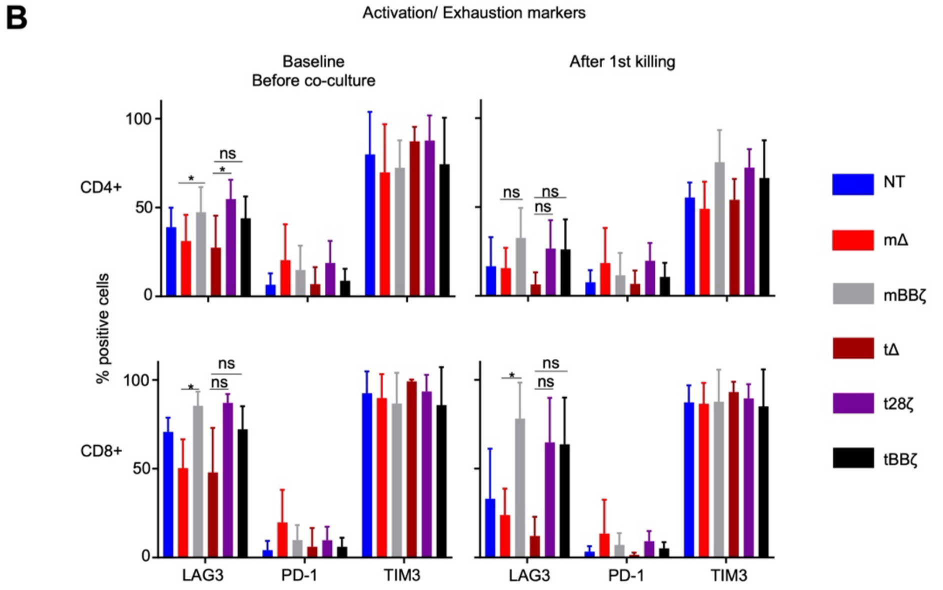

**Supplementary Figure 3. APRIL binding domain configuration and type of signaling endo-domains impact memory phenotype and activation/ exhaustion state of APRIL CAR T cells.** (**A**) Stacked bar graphs depict percent effector (E, blue), naive (N, red), effector memory (EM, grey) and central memory (CM, black) cells in NT or APRIL CAR T cells before (left) or after tumor challenge (right). T cells were gated on live cells, then on CD4 or CD8 positive cells, and finally on the CD271 positive fraction. n=6, mean±SD. One-way ANOVA on CM frequencies, significance levels: *p<0.05, **p<0.01, ***p<0.001, **** p<0.0001. Statistics on bar graphs compare percentage CM cells of mΔ control to each other CAR, the t28ζ vs tBBζ comparison is shown separately. (**B**) Bar graphs analyzing cell surface levels of activation/ exhaustion markers (LAG-3, PD-1, TIM-3) in APRIL CAR T cells before (left) and after (right) tumor challenge. n=6, mean±SD, One-way ANOVA, significance levels: *p<0.05.

### Supplementary Figure 4

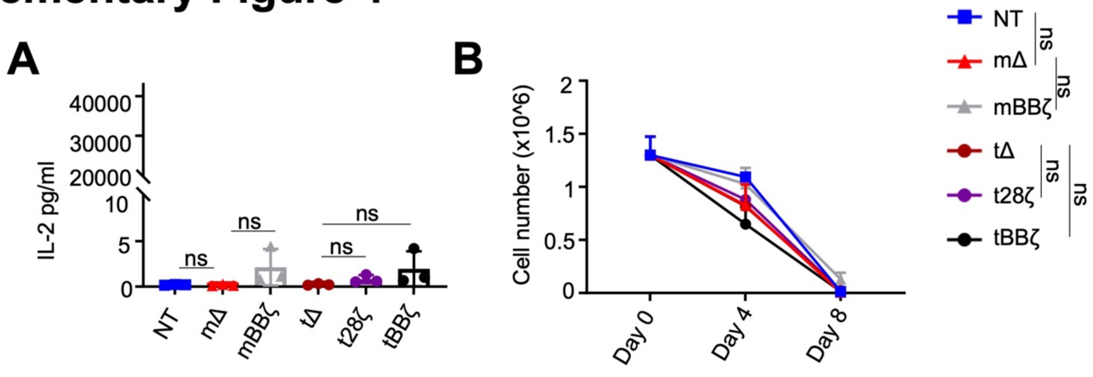


**Supplementary Figure 4. Assessment of tonic CAR activity.** (**A**) IL-2 production of CAR T cells over 24 hours in the absence of antigen challenge, n=3, mean±SD, one way ANOVA. (**B**) Basal CAR T cell expansion over time in absence target antigen exposure and exogenous cytokines, n=3, mean±SD, t-test on AUC. (**A, B**) ns: not significant.

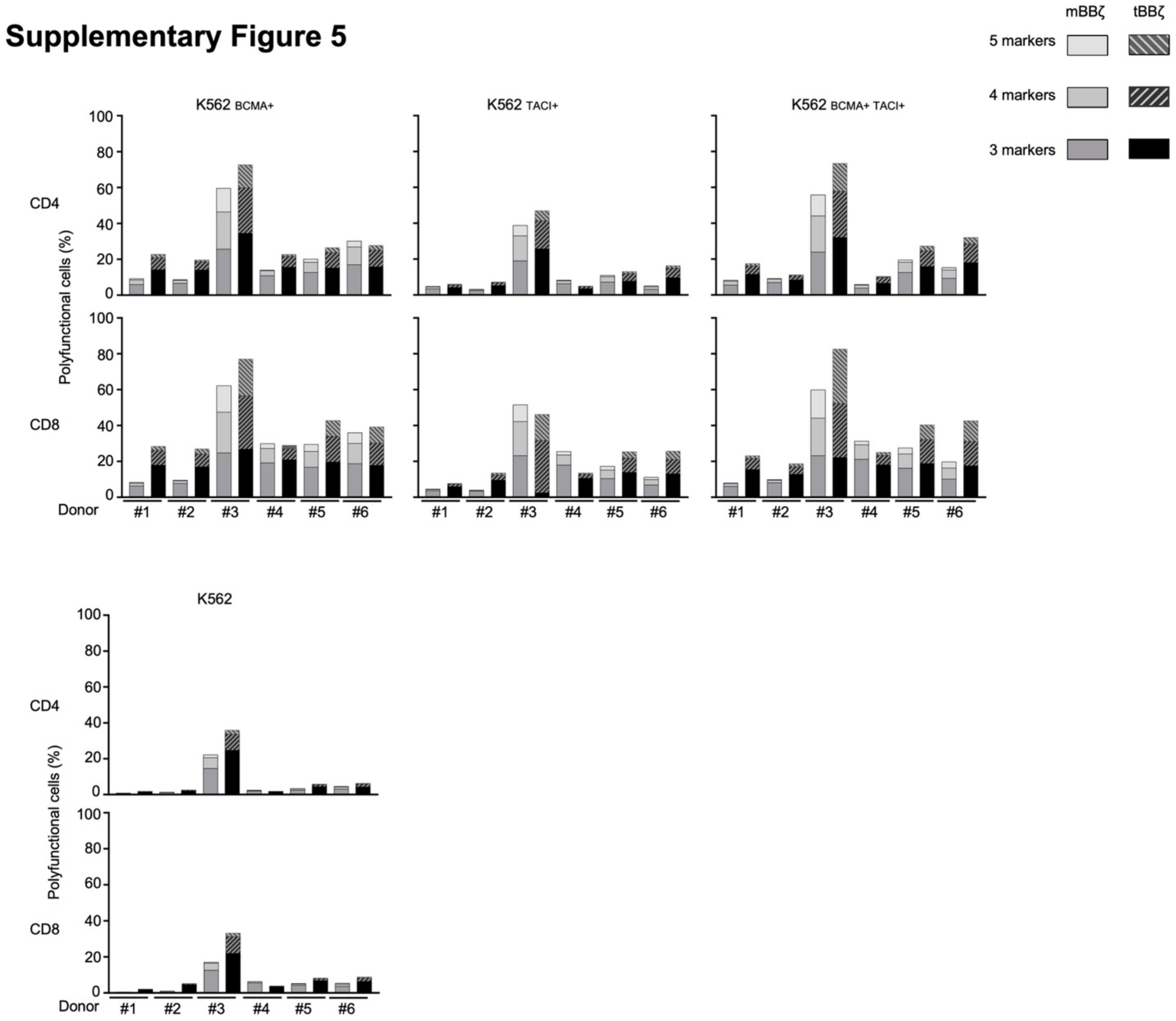


**Supplementary Figure 5. APRIL CAR T cell polyfunctionality in individual donors.** Stacked bar graphs depict the percentage of polyfunctional cells expressing at least 3, 4 or 5 markers, in individual donors comparing mBBζ to tBBζ challenged by tumor cells for 8 hours. Intracellular staining included TNF-α, IFN-γ, CD107a, GranzymeB and Perforin. n=6.

### Supplementary Figure 6

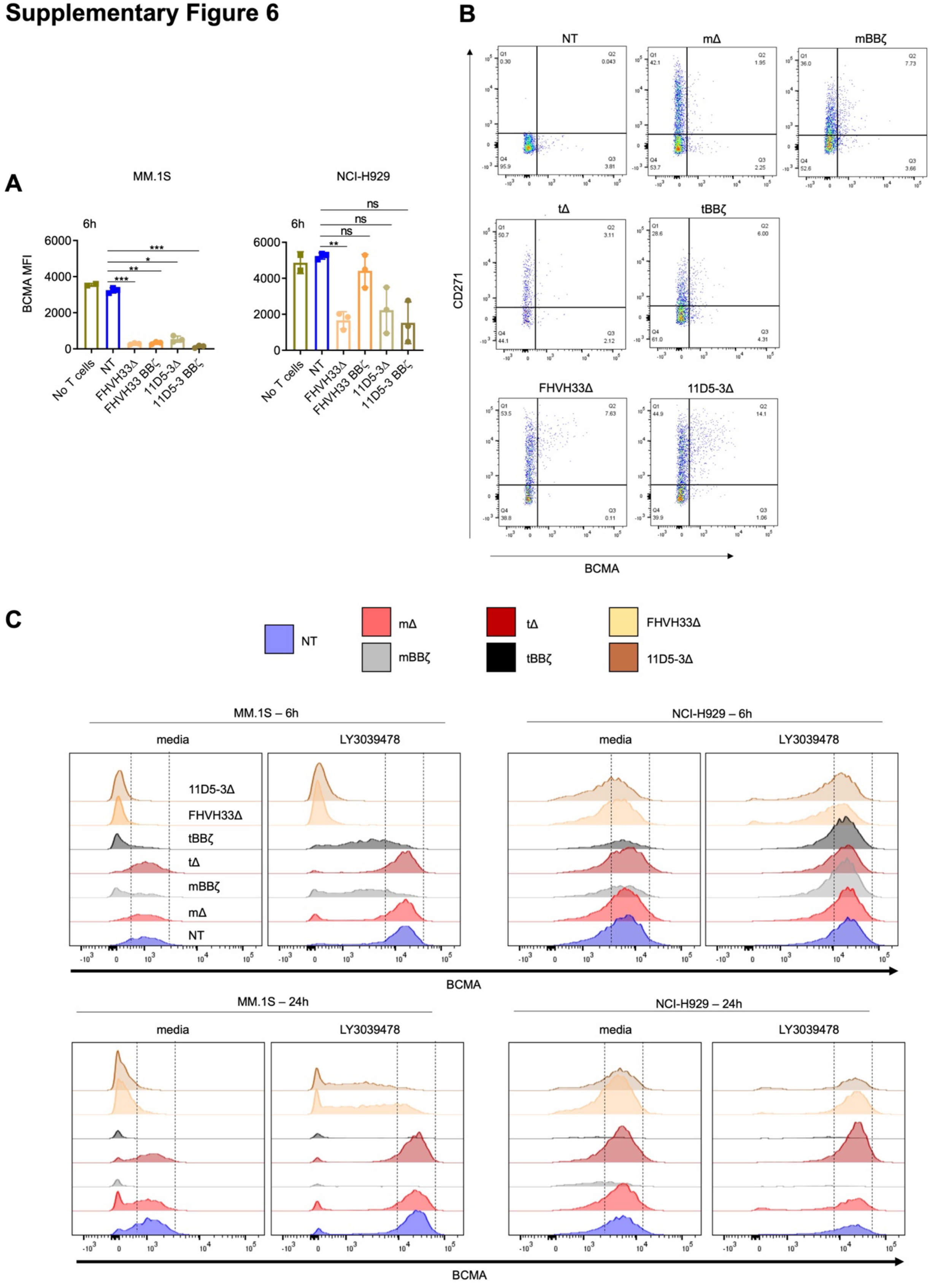

**Supplementary Figure 6. BCMA downmodulation on MM cell lines and trogocytosis by CAR T cells.** (**A**) Bar graphs show surface BCMA MFI from MM cells co-cultured with CAR T cells for 6 hours. n=3, mean±SD. One-way ANOVA, significance levels: *$p<0.05$, **$p<0.01$, ***$p<0.001$. (**B**) Representative flow cytometry dot plots showing BCMA detection on CAR T cells co-cultured with NCI-H929 cells for 6 hours. (**C**) Representative FACS histograms of surface BCMA staining on MM cells co-cultured with different CAR T cells from one donor for 6 or 24 hours, in presence or absence of $\gamma$-secretase inhibition (0.1µM LY3039478).

### Supplementary Figure 7

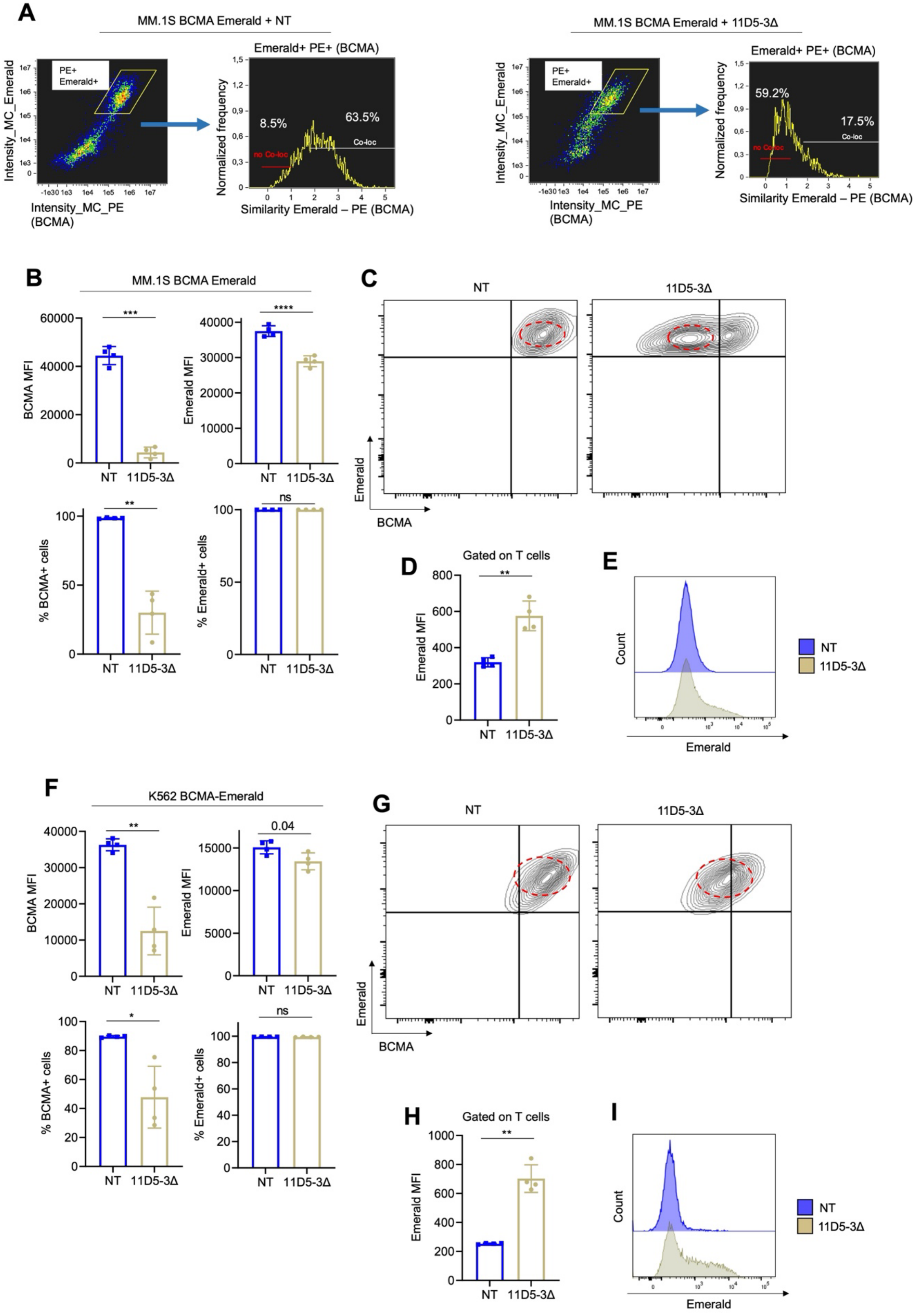

**Supplementary Figure 7. Evaluation of BCMA-Emerald internalization and trogocytosis upon co-culture with 11D5-3 scFv-based CAR T cells.** (**A**) MM.1S-BCMA-Emerald cells were co-cultured with NT or 11D5-3Δ CAR T cells for 1 hour. Membrane co-localization of BCMA-Emerald and cell surface BCMA (PE cell surface staining) was determined on the PE+ Emerald+ double positive population. Dot plots and histograms are representative of n=4 donors summarized in **Figure 4**. (**B**) Bar graphs show cell surface BCMA MFI and Emerald MFI (top) or percent cell surface BCMA+ or Emerald+ cells (bottom) after a 1-hour co-culture of T cells with MM.1S-BCMA-Emerald cells. (**C**) Representative FACS plot for cell surface BCMA and Emerald. (**D**) Emerald MFI on T cells and (**E**) representative histogram. (**F**) Bar graphs show cell surface BCMA MFI and Emerald MFI (top) or percent cell surface BCMA+ or Emerald+ cells (bottom) after a 1-hour co-culture of T cells with K562-BCMA-Emerald cells. (**G**) Representative FACS plot for cell surface BCMA and Emerald. (**H**) Emerald MFI on T cells and (**I**) representative histogram. Summary of n=4, mean±SD, paired t-test, significance levels: ns=not significant, *p<0.05, **p<0.01, ***p<0.001, ****p<0.0001.

### Supplementary Figure 8

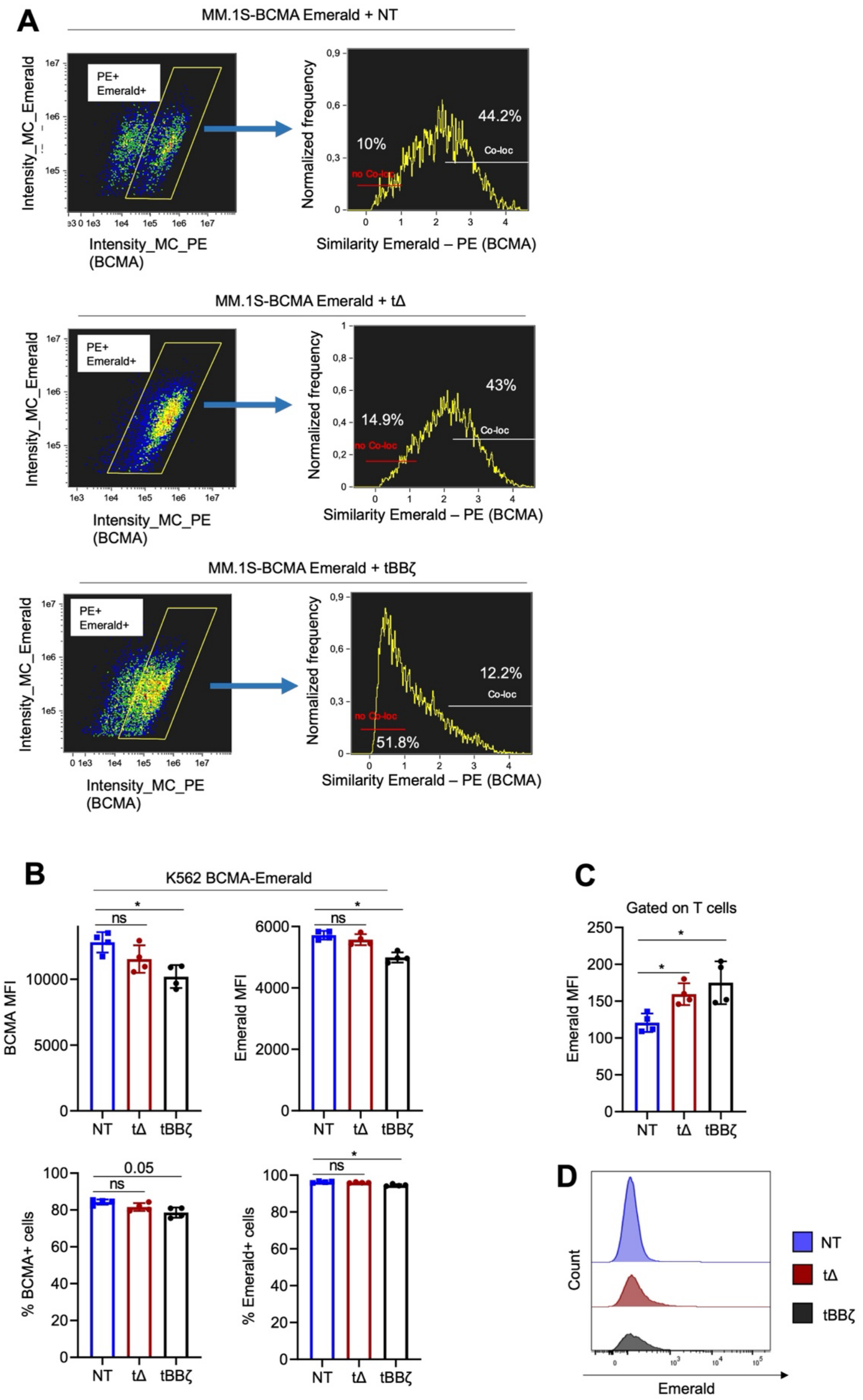

**Supplementary Figure 8. Evaluation of BCMA-Emerald internalization and trogocytosis upon co-culture with APRIL CAR T cells.** (**A**) MM.1S-BCMA-Emerald cells were co-cultured with NT, t$\Delta$ or tBB$\zeta$ CAR T cells for 1 hour. Membrane co-localization of BCMA-Emerald and cell surface BCMA (PE cell surface staining) was determined on the PE+ Emerald+ double positive population. Dot plots and histograms are representative of n=4 donors summarized in **Figure 4**. (**B**) K562-BCMA-Emerald cells were co-cultured with NT, t$\Delta$ or tBB$\zeta$ CAR T cells for 1 hour, bar graphs show BCMA and Emerald MFI or percentage of cell surface BCMA or Emerald positive cells. (**C**) Emerald MFI on T cells and (**D**) representative histogram. Summary of n=4, mean±SD, paired t-test, significance levels: ns=not significant, *p<0.05, **p<0.01, ***p<0.001, ****p<0.0001.

### Supplementary Figure 9

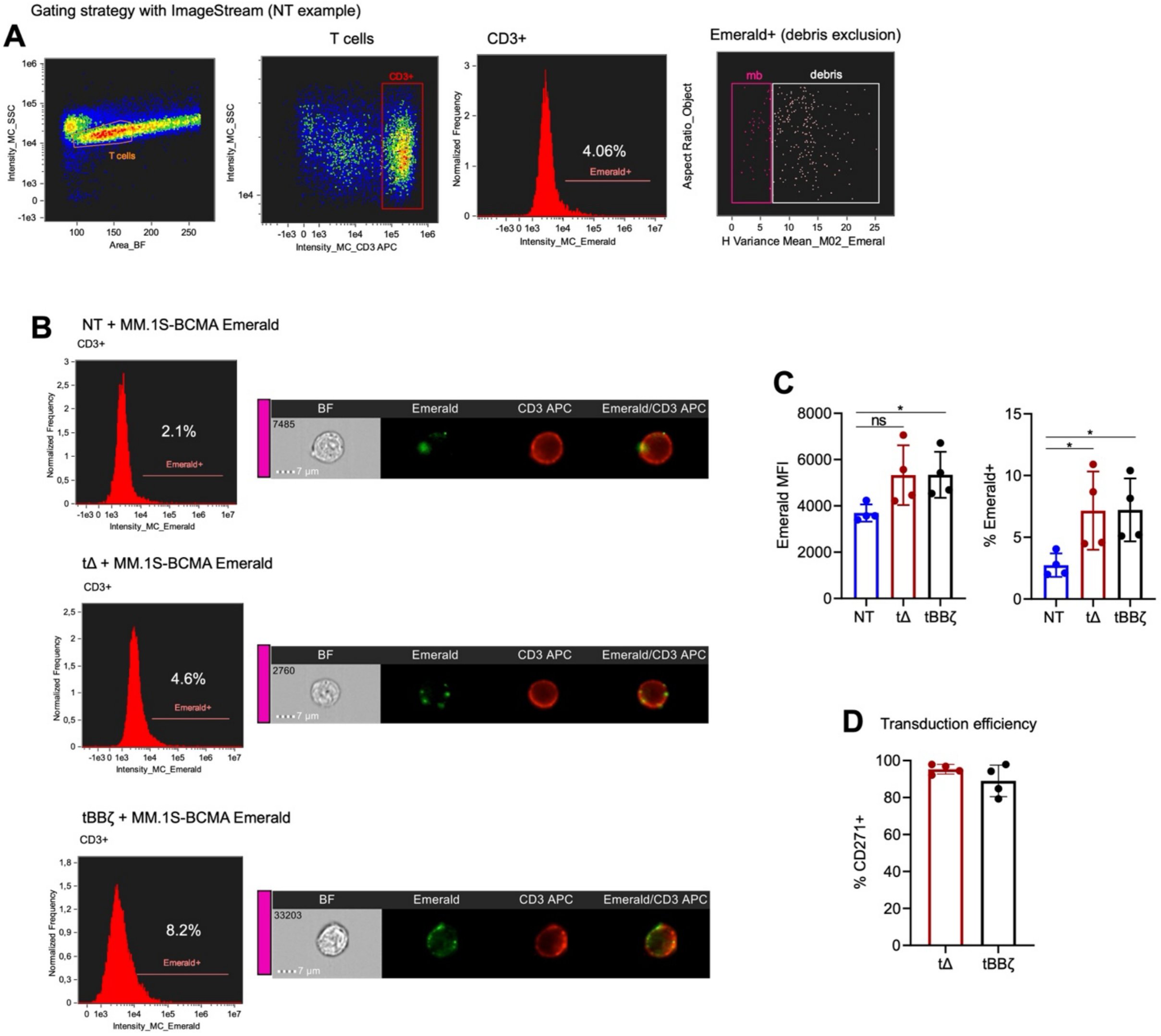


**Supplementary Figure 9. Image stream analysis reveals trogocytosis of BCMA-Emerald by APRIL CAR T cells.** (**A-C**) MM.1S-BCMA Emerald were co-cultured with NT, tΔ or tBBζ CAR T cells for 1h. (**A**) Gating strategy used to gate first on T cells, and then on Emerald+ population by excluding debris. (**B**) Representative graphs show percentage of Emerald+ T cells in NT, tΔ and tBBζ conditions, with representative images of Emerald and CD3 signals in the cell membrane. (**C**) Summary graphs show Emerald MFI on T cells (left) and percent Emerald positive T cells (right). (**D**) Bar graph depicting the percentage of CD271+ cells in tΔ and tBBζ conditions. (**C, D**) n=4, mean±SD, paired t-test, significance levels: ns=not significant, *p<0.05.

12. Henery S, George T, Hall B, et al. Quantitative image based apoptotic index measurement using multispectral imaging flow cytometry: a comparison with

standard photometric methods. *Apoptosis* 2008;13(8):1054-63. doi: 10.1007/s10495-008-0227-4 [published Online First: 2008/06/11]

13. George TC, Fanning SL, Fitzgerald-Bocarsly P, et al. Quantitative measurement of nuclear translocation events using similarity analysis of multispectral cellular images obtained in flow. *J Immunol Methods* 2006;311(1-2):117-29. doi: 10.1016/j.jim.2006.01.018 [published Online First: 2006/03/28]
14. Tischer C, Ravindran A, Reither S, et al. BigDataProcessor2: A free and open-source Fiji plugin for inspection and processing of TB sized image data. *Bioinformatics* 2021 doi: 10.1093/bioinformatics/btab106 [published Online First: 2021/02/18]
